# Predicting Pre-treatment Resistance or Post-treatment Effect? A Systematic Benchmarking of Single-Cell Drug Response Models

**DOI:** 10.64898/2026.04.10.717709

**Authors:** Li Shen, Xinliang Sun, Shuyu Zheng, Ali Hashmi, Johanna Eriksson, Harri Mustonen, Hanna Seppänen, Bairong Shen, Min Li, Markus Vähä-Koskela, Jing Tang

**Affiliations:** Research Program in Systems Oncology, Faculty of Medicine, University of Helsinki, Helsinki, 00014, Finland; School of Computer Science and Engineering, Central South University, Changsha, 410083, China; Translational Cancer Medicine Research Program, Faculty of Medicine, University of Helsinki, Helsinki, 00290, Finland; Department of Surgery, Faculty of Medicine, University of Helsinki and Helsinki University Hospital, Helsinki, 00290, Finland; iCAN Digital Precision Cancer Medicine Flagship, Helsinki, 00014, Finland; Department of Urology and Institutes for Systems Genetics, Frontiers Science Center for Disease-related Molecular Network, West China Hospital, Sichuan University, Chengdu, 610041, China; Institute for Molecular Medicine FIMM, Helsinki Institute for Life Science, University of Helsinki, Helsinki, 00014, Finland

**Author notes:** Corresponding author: Dr. Jing Tang.

**Keywords:** Single-cell RNA sequencing, Drug response prediction, Model benchmarking, Machine learning, Deep learning

## Abstract

Intratumoral heterogeneity drives variable drug responses in cancer. Single-cell RNA sequencing (scRNA-seq) enables characterization of such heterogeneity and prediction of drug response at single-cell resolution. Accordingly, various computational models have been developed to infer drug response from scRNA-seq data. However, their performance, robustness, and generalizability across different biological contexts remain insufficiently evaluated.

To address this gap, we benchmarked representative single-cell drug response prediction models using 26 curated datasets comprising over 760,000 cells across 12 cancer types and 21 therapeutic agents. We constructed balanced and imbalanced scenarios to explore model behavior under different label distributions. To address the lack of ground-truth labels in conventional scRNA-seq datasets, we incorporated two lineage-tracing datasets with experimentally validated drug-response annotations, enabling evaluation in a clinically relevant pre-treatment prediction setting.

Our results show that prediction performance was markedly higher in cell lines than in tissue samples and varied substantially across datasets, treatment types, and imbalance levels. Under increasing class imbalance, several methods showed reduced performance, and no single method consistently dominated across all evaluation settings. In lineage-tracing datasets, treatment-associated states were more readily distinguished than lineage-defined pre-sensitive and pre-resistant states, while transcriptomic analyses indicated that pre-existing resistance-related signals were detectable but relatively subtle. For scDEAL, which showed competitive performance across several benchmark settings, label-substitution and ablation analyses further indicated that its performance was jointly influenced by training-label definition and architectural design. Application to an independent PDAC organoid dataset showed that scDEAL predictions were associated with treatment-related transcriptional state changes and biological programs.

In summary, our study systematically characterizes the performance and limitations of current single-cell drug response prediction approaches across diverse evaluation settings. These results highlight the influence of dataset characteristics, class imbalance, training design, and the strength of resistance-associated signals on model performance, providing a reference for the evaluation and further development of single-cell drug response prediction methods.

## Introduction

Drug resistance remains a major challenge in effective cancer treatment due to the highly heterogeneous responses of tumor cells to therapeutic agents [1]. Traditional Bulk RNA-seq methods aggregate gene expression across millions of cells, which often obscure rare subpopulations responsible for cellular heterogeneity in drug responses [2]. In contrast, single-cell RNA sequencing (scRNA-seq) enables high-resolution profiling at the individual cell level, allowing researchers to monitor treatment-induced transcriptional changes, identify resistant subpopulations, and decipher the drug response heterogeneity. With the rapid accumulation of single-cell datasets generated under diverse treatment conditions, an increasing number of computational methods have been developed to predict the drug response at single-cell resolution [3]. These methods have significantly enhanced our ability to identify drug-resistant subpopulations and optimize combinatorial therapies, further advancing the discovery of personalized treatment strategies [4].

Recently, large foundation models pretrained on massive single-cell datasets have shown strong performance across diverse analysis tasks [5, 6]. For example, the scDrugMap framework represents a significant step forward by benchmarking multiple single-cell foundation models (scFMs) for drug response prediction [7]. Nevertheless, scFMs often require substantial computational resources and long runtimes, which can be prohibitive for users with limited computational infrastructure [8, 9]. In contrast, traditional machine learning methods are generally computationally efficient, making them more accessible to biomedical researchers and clinicians without extensive AI expertise [10].

Despite these achievements, current methods remain constrained by several key limitations affecting their robustness and broad applicability. Many methods were developed using limited data sources, such as the Genomics of Drug Sensitivity in Cancer (GDSC) [11] and the Cancer Cell Line Encyclopedia (CCLE) [12], raising concerns about their generalizability across diverse biological contexts [13, 14].

Moreover, real-world single-cell datasets often exhibit substantial class imbalance, where the proportions of drug-resistant and drug-sensitive cells may differ significantly across samples. Such data imbalance can affect both model training and evaluation [15, 16], yet its impact has not been systematically assessed. On the other hand, existing methods are typically evaluated using different datasets and validation strategies. The lack of standardized evaluation study creates uncertainty for translational researchers when selecting appropriate methods for their own datasets [17]. Therefore, performing a systematic and transparent benchmarking analysis is essential for guiding method selection in line with the FAIR (Findable, Accessible, Interoperable, Reusable) principles.

Furthermore, to establish a more clinically relevant benchmarking setting, where patient samples are collected prior to treatment and the task is to predict which cell populations are intrinsically resistant, we note that conventional scRNA-seq datasets containing only pre- and post-treatment samples are insufficient to provide reliable ground truth labels [18]. In many commonly used scRNA-seq datasets, drug-sensitive cells are typically assumed to correspond to pre-treatment samples or samples from responding patients, whereas resistant cells are assumed to correspond to post-treatment samples or non-responding patients. However, these assumptions lack definitive single-cell validation, as drug response labels are typically inferred from treatment status rather than experimentally validated at the single-cell level. Consequently, evaluating drug response prediction models under such proxy labelling schemes may introduce bias and lead to misleading performance estimates, potentially limiting their reliability for clinical translation [19].

In this work, we conducted a systematic benchmarking study to assess key aspects of single-cell drug response prediction, including classification accuracy, computational efficiency, and cross-dataset generalizability. To address real-world application needs, we considered class balance and imbalance as distinct scenarios. Furthermore, to overcome the limitation of ground-truth labels, we included two independent lineage-tracing datasets, ReSisTrace [20] and Watermelon [21], where pre-treatment cells can be linked to subsequent treatment outcomes through lineage information. This enables pre-treatment cells to be classified as pre-sensitive or pre-resistant according to the subsequent fate of their corresponding lineages, providing experimentally supported response labels for evaluating pre-treatment resistance prediction. Our results reveal that while certain methods achieve promising performance under specific conditions, most exhibit varying degrees of instability, where notable performance fluctuations across datasets are frequently observed even for the same drug treatment predicted by the best-performing models. Lineage-tracing evaluations further showed that pre-existing resistance was more difficult to predict than treatment-associated states.

Given the relatively strong performance of scDEAL across several benchmark settings, we further examined its performance determinants through training-label substitution and ablation analyses, and evaluated the biological relevance of its predictions in an in-house PDAC organoid dataset. Taken together, our benchmarking analysis provides a valuable reference for guiding the evaluation and improvement of computational methods in predicting drug response at single-cell resolution. Our findings highlight the need for more robust and clinically relevant approaches to better leverage the single-cell data for the understanding of drug response.

## Materials and Methods

### Data collection and preprocessing

We obtained single-cell datasets with annotated drug response labels from the DRMREF (https://ccsm.uth.edu/DRMref/) database [22], which integrates 42 single-cell drug response datasets, among which 26 single-drug treatment datasets were selected for benchmarking. The selected datasets cover 12 cancer types and 19 molecular drugs as well as two cellular therapies, comprising a total of 760,090 cells. For each dataset, a binary label (resistant vs. sensitive) was used to annotate the drug-response status of each cell for a given treatment. In cell line datasets, cells from pre-treatment samples were labeled as sensitive and post-treatment cells as resistant. In patient-derived datasets, cells from non-responding patients were labeled as resistant, whereas cells from responding patients were labeled as sensitive.

To ensure consistency across datasets, we performed standardized preprocessing of cell-level metadata, including converting condition labels to lowercase, removing whitespace, and correcting common spelling errors. Uniform quality control criteria were then applied across all datasets: cells with fewer than 1,000 detected genes or with a mitochondrial read fraction greater than 15% were removed, and genes expressed in fewer than three cells were discarded. Detailed dataset characteristics are summarized in **Table 1**.

**Table 1.** Summary of the single-cell datasets used in the study.

| **Dataset** | **Treatment** | **Mechanisms of action (MoA)** | **Sample type** | **Disease** | **Total cells** | **Resistant** | **Sensitive** |
| --- | --- | --- | --- | --- | --- | --- | --- |
| GSE104987 | KDM5-C70 | Epigenetic inhibitor targeting H3K4 demethylation | MCF7 cell line | Breast cancer | 2669 | 1597 | 1072 |
| GSE108397 | PLX-4720 | Selective inhibition of mutant BRAF in the MAPK pathway | 451Lu cell line | Melanoma | 6548 | 3262 | 3286 |
| GSE111014 | Ibrutinib | BTK inhibitor suppressing BCR signaling | Peripheral blood mononuclear cells | Chronic lymphocytic leukemia | 24790 | 12645 | 12145 |
| GSE115251 | TAE684 | ALK tyrosine kinase inhibitor | Kelly cell line | Neuroblastoma | 11790 | 6298 | 5492 |
| GSE131984_JQ1 | JQ1 | BET bromodomain inhibitor suppressing transcriptional programs | SUM159 cell line | Breast cancer | 2167 | 1565 | 602 |
| GSE131984_Pac | Paclitaxel | Microtubule-stabilizing agent causing mitotic arrest | SUM159 cell line | Breast cancer | 1354 | 752 | 602 |
| GSE131984_Pal | Palbociclib | CDK4/6 inhibitor inducing G1 cell-cycle arrest | SUM159 cell line | Breast cancer | 1295 | 693 | 602 |
| GSE139386 | Ceritinib | ALK tyrosine kinase inhibitor | H3122 cell line | Lung cancer | 6986 | 4622 | 2364 |
| GSE140440 | Docetaxel | Microtubule depolymerization inhibitor | DU145 and PC cell lines | Prostate cancer | 284 | 150 | 134 |
| GSE149214 | Erlotinib | EGFR tyrosine kinase inhibitor | PC9 cell line | Lung cancer | 2393 | 1282 | 1111 |
| GSE152469 | Ibrutinib | BTK inhibitor suppressing BCR signaling | Peripheral blood mononuclear cells | Chronic lymphocytic leukemia | 9264 | 4932 | 4332 |
| GSE153697 | anti-CD19 CAR-T | CD19-directed CAR-engineered T cells inducing antigen-specific cytotoxicity | Bone marrow aspirate | Acute lymphoblastic leukemia | 2919 | 1206 | 1713 |
| GSE158457 | Dasatinib | Multi-kinase tyrosine kinase inhibitor | PDX cells | Acute lymphoblastic leukemia | 26026 | 14197 | 11829 |
| GSE162117 | Ficlatuzumab | Neutralizing antibody against HGF | Peripheral blood mononuclear cells | Acute myeloid leukemia | 43486 | 17026 | 26460 |
| GSE163836 | Paclitaxel | Microtubule-stabilizing agent causing mitotic arrest | FCIBC02 cell line | Breast cancer | 7358 | 5380 | 1978 |
| GSE164237 | Pembrolizumab | Immune checkpoint inhibitor blocking PD-1 signaling | Peripheral blood mononuclear cells | Melanoma | 23276 | 11634 | 11642 |
| GSE164551 | anti-BCMA CAR-T | BCMA-directed CAR-engineered T cells inducing antigen-specific cytotoxicity | Bone marrow aspirate | Multiple myeloma | 7736 | 3075 | 4661 |
| GSE164897_Vem | Vemurafenib | BRAF inhibitor blocking MAPK signaling | A375 cell line | Melanoma | 11008 | 4322 | 6686 |
| GSE168668 | Enzalutamide | Androgen receptor antagonist inhibiting AR signaling | LNCaP cell line | Prostate cancer | 10698 | 8542 | 2156 |
| GSE169246_PacB lood | Paclitaxel | Microtubule-stabilizing agent causing mitotic arrest | Peripheral blood mononuclear cells | Breast cancer | 120823 | 61803 | 59020 |
| GSE169246_PacT issue | Paclitaxel | Microtubule-stabilizing agent causing mitotic arrest | Tumor tissue | Breast cancer | 67140 | 36913 | 30227 |
| GSE175716 | Sorafenib | Multi-kinase inhibitor targeting proliferation and angiogenesis | PDX | Liver cancer | 6823 | 958 | 5865 |
| GSE186960 | Gemcitabine | Nucleoside analog inhibiting DNA replication | Panc1 cell line | Pancreatic cancer | 6136 | 5438 | 698 |
| GSE199333 | Gilteritinib | FLT3 inhibitor blocking oncogenic kinase signaling | Bone marrow aspirate | Acute myeloid leukemia | 105004 | 54241 | 50763 |
| GSE212217 | Pembrolizumab | Immune checkpoint inhibitor blocking PD-1 signaling | Peripheral blood mononuclear cells | Endometrial cancer | 224785 | 93974 | 130811 |
| PMID34715028 | Nivolumab | Immune checkpoint inhibitor blocking PD-1 signaling | Tumor tissue | Kidney cancer | 27332 | 8610 | 18722 |

### Balanced data generation

To provide a baseline scenario without class imbalance, we generated balanced data subsets of fixed sample sizes from the original datasets. Balanced data subsets were constructed at five predefined sample sizes: 500, 1000, 2000, 3000, and 5000 cells.

For each sample size *k*, we required that each drug response label contains at least 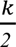 cells. For datasets fulfilling the requirement, a stratified random sampling was then performed to extract 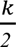 cells from each drug response label, forming a balanced subset containing a total of *k* cells. Sampling was performed without replacement, and repeated three times using fixed random seeds to ensure reproducibility.

### Imbalanced data generation

In addition to the balanced sampling scenario, we also constructed imbalanced subsets to evaluate model performance under different levels of class imbalance. Data subsets were generated under four predefined minority-to-majority ratios including: 1:3, 1:10, 1:30, and 1:100. For each ratio, the numbers of minority and majority cells were determined by imposing an upper limit of 10,000 cells per subset:

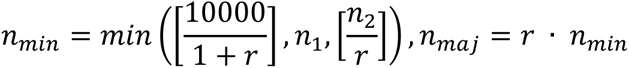

Here, *n*_1_ and *n*_2_ denote the available numbers of minority and majority cells in the original dataset, respectively; *r* is the target imbalance ratio (majority versus minority). Within each ratio setting, stratified random sampling was performed independently for the minority and majority classes to obtain *n_min_* and *n_maj_* cells, respectively. Sampling was repeated without replacement three times with fixed random seeds.

### Overview of the benchmarking methods

We benchmarked nine representative single-cell drug response prediction methods, including Beyondcell [23], CaDRReS-sc [24], DREEP [25], DrugFormer [26], Precily [27], SCAD [28], scDEAL [29], scDr [30], and scIDUC [31]. These methods were selected to represent a diverse range of modeling strategies, ranging from classical statistical learning to deep learning–based models. In addition, we evaluated two recent single-cell foundation models, scGPT [6] and scFoundation [5], as complementary benchmarks in the balanced, imbalanced, and lineage-tracing settings. To ensure a fair comparison, each method was implemented following the procedures described in their original publications or GitHub repositories, without additional modifications. The implementation workflows, including input preparation and parameter settings, are available in the Supplementary Methods. The evaluated methods use different modeling strategies and output definitions, whereas their outputs provide quantitative estimates of cell-level drug response, expressed as probabilities or response scores. The interpretation of these outputs followed the definition of each original method. We assessed their predictions against the benchmark response labels using a common set of evaluation metrics.

### Lineage tracing datasets

To obtain reliable single-cell–level ground truth for drug response prediction, we included two independent lineage-tracing datasets, ReSisTrace [20] and Watermelon [21]. With ReSisTrace, cancer cells are labeled with unique 20-bp random barcodes, synchronized, and allowed to undergo one cell division. Sister cells sharing the same barcode are then separated into two groups: one group undergoes single-cell RNA sequencing prior to treatment, while the other is exposed to drug treatment followed by sequencing of surviving cells. By matching barcode identities between pre- and post-treatment samples, ReSisTrace enables direct classification of pre-treatment cells as pre-resistant or pre-sensitive, thereby providing truly cell-resolved response labels rather than population- or clone-level approximations.

For ReSisTrace, we analyzed the Kuramochi ovarian cancer datasets profiled under Olaparib treatments. The scRNA-seq data from both before- and after-treatment samples, together with the corresponding barcode-based lineage annotations, were processed following the procedures described in the original publication [20]. To address class imbalance, we also performed three independent random samplings in which a subset of pre-sensitive cells equal in size to the pre-resistant set was drawn without replacement.

The Watermelon dataset (GSE150949) provided an independent lineage-tracing system using PC9 lung cancer cells sampled before treatment (D0) and at D3, D7, and D14 after treatment. Only cells assigned to a single lineage barcode were used for response-state definition. Lineages detected at D0 and retained at D3, D7, and D14 were classified as resistant lineages, and their D0 cells were labeled as pre-resistant.

Lineages detected at D0 but absent from all three post-treatment time points were classified as sensitive lineages, and their D0 cells were labeled as pre-sensitive. Post-treatment cells belonging to resistant lineages were labeled as resistant. For intrinsic resistance prediction, D0 pre-sensitive and pre-resistant cells were compared. To reduce class imbalance, three independent balanced subsets were generated by retaining all cells from the minority class and randomly subsampling an equal number of cells from the majority class. For comparison with the conventional pre- versus post-treatment setting, D0 pre-sensitive cells and D3 resistant cells were used.

Bioinformatics analyses were also conducted to characterize transcriptomic differences between pre-sensitive and pre-resistant cells in both lineage-tracing datasets. Detailed procedures are provided in the Supplementary Methods.

### Evaluation metrics

Model performance was evaluated using AUROC, normalized AUPRC, accuracy, and F1-score across all scenarios, with normalized AUPRC providing primary insight under imbalanced settings. For methods requiring model training, metrics were averaged via five-fold cross-validation, whereas for off-the-shelf tools (e.g., R packages with direct prediction functions), performance was reported based on their single-run prediction results. The normalized AUPRC is defined as:

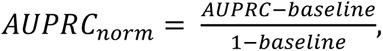

where *baseline* corresponds to the proportion of positive samples in the test set, representing the expected AUPRC of a random classifier. By adjusting for this baseline, the normalized AUPRC provides a scale-invariant metric that allows fair comparison across datasets with different levels of class imbalance. Under this definition, a value of 0 corresponds to random prediction, 1 indicates perfect classification, and negative values suggest performance worse than random prediction.

To reduce the influence of differences in baseline dataset difficulty when comparing models, we also performed within-dataset comparisons. For each dataset and sampling setting, repeated measurements for each method were first averaged. Performance was then centered by subtracting the median performance across available methods within the same dataset and setting. We also calculated within-dataset ranks and normalized them to a 0–1 scale, where 0 represents the best-performing method and 1 the lowest-performing method within the corresponding dataset and setting. AUROC was used for balanced datasets and normalized AUPRC for imbalanced datasets.

### Training label substitution experiment

The training labels used by scDEAL and SCAD were first compared using their shared drugs and cell lines. Drug names and cell-line identifiers were standardized across the two label sets, and drug–cell line pairs with available labels from both methods were retained. Drugs represented by at least 30 common cell lines were included in the comparison. For each drug, we calculated the fraction of resistant cell lines under each label definition and measured label agreement using Cohen’s kappa. The common cell lines were further divided into four groups according to their scDEAL/SCAD label combinations: S/S, S/R, R/S, and R/R, where S and R represents sensitive and resistant, respectively. GDSC logIC50 values were standardized within each drug before comparison across the four groups.

We next performed training label substitution between scDEAL and SCAD using dataset–drug pairs available for both methods. The experiment was conducted on the previously generated balanced datasets and the 1:3 and 1:30 imbalanced datasets using the same three sampling rounds. For each method, models were evaluated using either their original training labels or the labels from the other method. For SCAD, source cell lines with available labels under both label definitions were retained, and the same source expression data, data splits, and model configuration were used for the two label conditions. For scDEAL, the model configuration selected from the original benchmark was retained when the training labels were replaced. In the balanced setting, model configurations were selected according to AUROC, whereas normalized AUPRC was used for the imbalanced settings. Performance was evaluated using accuracy, AUROC, F1 score, and normalized AUPRC for the imbalanced datasets. For SCAD, performance was first calculated for each cross-validation split and then averaged across the five splits.

To further separate the effects of training labels and model architecture, four label– architecture combinations were defined: SCAD architecture with SCAD labels (A), SCAD architecture with scDEAL labels (B), scDEAL architecture with SCAD labels (C), and scDEAL architecture with scDEAL labels (D). Performance was first averaged across the three sampling rounds for each dataset–drug pair. The scDEAL label effect was calculated as D − C, the scDEAL architecture effect as D − B, the SCAD label effect as A − B, and the SCAD architecture effect as A − C. The label– architecture interaction was calculated as (D − C) − (B − A). Mean effect sizes and 95% confidence intervals were estimated using 5,000 bootstrap resamples.

### scDEAL ablation analysis

To evaluate the contribution of different components of scDEAL, we performed ablation analysis using the same dataset–drug pairs as in the training label substitution experiment. The analysis was performed on the 1,000-cell balanced datasets and the 1:3 and 1:30 imbalanced datasets. Four components were evaluated: transfer learning, cluster regularization, DAE pretraining, and source-data integration. For the transfer-learning ablation, predictions before the transfer-learning step were used. Cluster regularization was removed from the original model, while DAE pretraining was replaced with a standard autoencoder (AE). Source-data integration was evaluated by comparing the full model trained on integrated GDSC and CCLE data with models trained on GDSC or CCLE alone. The bottleneck size and sampling method selected for the full scDEAL model were retained for the corresponding ablation experiments.

For each dataset, sampling round, and evaluation setting, the contribution of each component was calculated as:

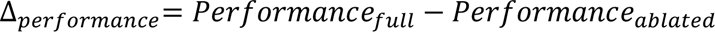

For source-data integration, the better-performing single-source model was used as the ablated reference. Components were ranked according to their mean Δ*_performance_* within each evaluation condition, with rank 1 indicating the largest contribution. The contribution of training-label definition, calculated from the training label substitution experiment, was also included in the overall ranking. For balanced datasets, AUROC, F1 score, and accuracy were used for ranking, while normalized AUPRC, AUROC, and F1 score were used for the imbalanced datasets. The mean rank across all evaluation conditions was then calculated to summarize the overall contribution of each model component.

### PDAC organoid dataset and downstream analysis

PDAC organoid lines PO83T and PO84T were established and characterized as previously described [32] and cultured in Matrigel following the protocol of Boj et al. [33]. For transcriptional profiling of drug responses, organoids were treated with 12.5 nM trametinib or 500 nM paclitaxel for 72 h, with DMSO used as the vehicle control. Single-cell RNA-seq libraries were generated using the 10x Genomics Chromium platform, and the resulting data were processed using Seurat (version 4.3.0) [34]. Highly variable genes were identified for each sample, and batch effects across samples were corrected using Harmony based on the first 30 principal components [35].

For downstream characterization of scDEAL predictions, cells were stratified according to the median predicted drug-sensitivity probability. Cell-cycle effects were regressed out before differential expression analysis. Pathway activities were evaluated using single-sample GSEA implemented in GSVA [36], while custom drug-response and pathway signatures were scored using AUCell [37]. Pseudotime trajectories were reconstructed using Monocle 2 with DDRTree dimensionality reduction [38]. The top 300 and 1,000 differentially expressed genes ranked by absolute log2 fold change were used to order cells for the paclitaxel and trametinib datasets, respectively. Genes showing significant expression changes across the inferred trajectories were subsequently clustered and subjected to functional enrichment analysis using ClusterGVis [39]. Detailed procedures for organoid culture, single-cell dissociation, cell hashing, library preparation, and downstream analyses are provided in the Supplementary Methods.

### Statistical analysis

All statistical analyses and visualizations were performed in R (version 4.4.0, RRID:SCR_001905). Comparisons between two independent groups were performed using the Wilcoxon rank-sum test, while paired comparisons were performed using the Wilcoxon signed-rank test. For the label and architecture effect analysis, 95% confidence intervals were estimated using 5,000 bootstrap resamples. For the scDEAL ablation analysis, mean performance changes and their 95% confidence intervals were calculated across dataset–round combinations. Where multiple testing correction was applied, p-values were adjusted using the Benjamini–Hochberg method. A p-value or adjusted p-value < 0.05 was considered statistically significant unless otherwise specified.

## Results

### Benchmarking design and characteristics of the datasets

To conduct a systematic benchmarking analysis, we obtained 26 single-cell drug response datasets. The evaluation datasets were then organized into balanced and imbalanced scenarios. Across these scenarios, we evaluated nine computational models and compared their performance using a comprehensive set of classification metrics. In addition, the lineage-tracing datasets were used as ground truth references for more rigorous comparisons. **Figure 1A** provides the lineage tracing scheme and the analysis workflow.

**Figure 1.**
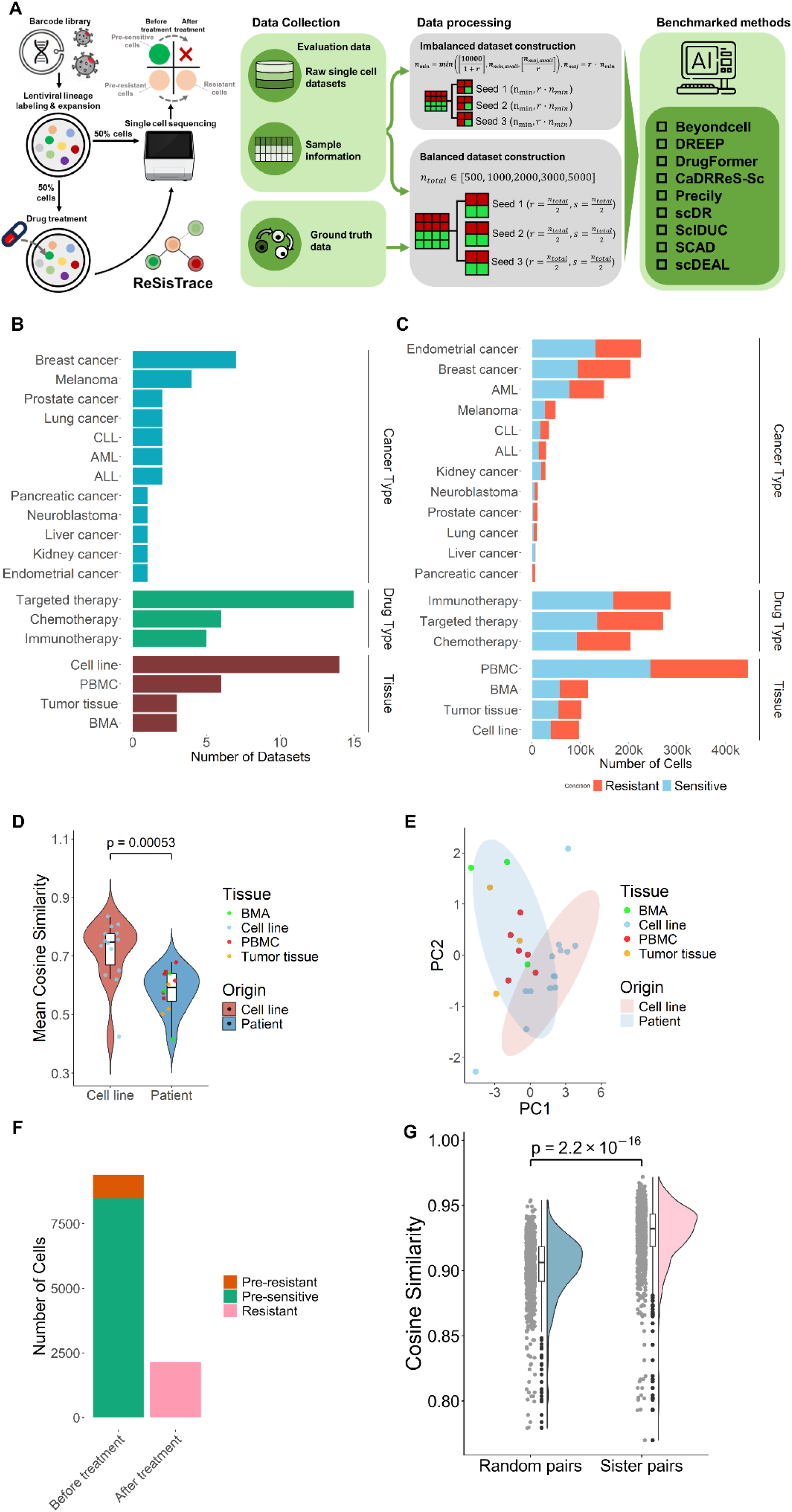
(A) Overview of the ground-truth data generation based on lineage tracing (left) and the benchmarking workflow (right). With the ReSisTrace lineage tracing, cancer cells are uniquely labelled by using lentiviral constructs, and are permitted to undergo a single doubling, after which half of the cells undergo scRNA-sequencing, while the remaining half are treated with anti-cancer therapy. The treated cells are then analyzed with scRNA-sequencing to identify the lineages of pre-resistant cells. Such a process allows a high-resolution analysis of single-cell drug responses in a pre-treatment sample. The benchmarking workflow utilized manually curated conventional single-cell drug response and lineage-tracing datasets. Following quality control, the conventional datasets were partitioned into balanced and imbalanced scenarios, while lineage-tracing datasets were also sampled into balanced splits to ensure an unbiased evaluation. Finally, nine single-cell drug response prediction methods were systematically assessed using scenario-appropriate performance metrics to enable a comprehensive comparison of model performance. **(B-C)** Summary of the evaluation datasets including the number of datasets **(B)** and number of cells **(C)** across tissue types, drug types, and cancer types. **(D-E)** Similarity between resistant and sensitive cells in the evaluation datasets. **(D)** Mean cosine similarity grouped by sample origin. **(E)** Sample separation based on principle components of aggregated gene expression profiles. **(F**-**G)** Characterization of the lineage-tracing dataset. **(F**) Distribution of cells across treatment time points, grouped by drug response labels. (**G**) Cosine similarity between sister cell pairs and random pairs. ALL: Acute lymphoblastic leukemia, AML: Acute myeloid leukemia, CLL: Chronic lymphocytic leukemia, PBMC: Peripheral blood mononuclear cells, BMA: Bone marrow aspirate.

The evaluation datasets cover 12 cancer types, including seven breast cancer datasets, four myeloma datasets, and two lung cancer datasets. Three major therapeutic modalities are well represented, including 15 targeted therapies, six chemotherapies, and five immunotherapies. The samples cover four specimen types including 14 cell lines, six peripheral blood mononuclear cell (PBMC) samples, three tumor tissues, and three bone marrow aspirates. Detailed descriptions are provided in **Table 1** and shown in **Figure 1B and C**. To further characterize transcriptional differences between drug-resistant and drug-sensitive cells, we computed the average cosine similarity between them within each dataset (**Figure 1D**). Cell line datasets showed significantly higher similarity compared with patient tissue datasets (Wilcoxon rank-sum test, p = 0.00053). PCA of aggregated expression profiles also revealed that cell line and patient tissue datasets formed two distinct clusters (**Figure 1E**). These results are consistent with the greater transcriptional heterogeneity observed in clinical samples [40]. The distribution of drug response labels in the lineage-tracing dataset is summarized in **Figure 1F**. Specifically, in the pre-treatment group, the population consisted of 8490 pre-sensitive cells and 904 pre-resistant cells, whereas the after-treatment group contained 2152 resistant cells. We further compared transcriptional similarity between annotated sister-cell pairs and randomly selected non-sister pairs. Sister-cell pairs showed substantially higher cosine similarity (Wilcoxon rank-sum test, p = 2.2 × 10⁻¹⁶) (**Figure 1G**). This high concordance confirms that the transcriptome of one sister cell serves as a reliable reference for the other, enabling the identification of primed resistance states across split samples [20].

### Performance on the Balanced Datasets

Considering that dataset sizes may influence the behavior of single-cell models [41], we performed subsampling ranging from 500 to 5000 cells for each dataset to examine how cell number affects prediction performance. As shown in **Figure 2A**, scDEAL achieved the highest median AUROC and ACC across all subsampling scenarios, with median AUROC values ranging from 0.67 to 0.82 and median ACC values ranging from 0.6 to 0.65. Mean performance trends further showed that increasing cell number did not consistently improve prediction performance: scDEAL and CaDRReS-sc showed fluctuating AUROC, whereas DREEP, Precily, and scDr generally declined with increasing cell number; scDEAL ACC showed a modest increase (**Figure S1A and B**). We compared the AUROC and ACC differences between scDEAL and other methods under different sample size settings (**Figure S2**). In small-sample scenarios (e.g., 500 or 1000 cells), scDEAL generally showed higher performance than most methods (Wilcoxon rank-sum test, p < 0.05). However, as the cell number increased to 2000 or more, although scDEAL remained significantly superior to Precily and scDr (Wilcoxon rank-sum test, p < 0.05), its statistical advantage over CaDRReS-sc and SCAD gradually reduced. On the other hand, scDEAL exhibited substantial dataset-level variability, with AUROC values ranging from 0.46 to 0.99 and ACC values from 0.17 to 0.97 across datasets. A similar pattern was also observed for CaDRReS-sc, whose AUROC and ACC values spanned 0.48-0.97 and 0.08-0.92, respectively. These wide performance ranges indicate substantial dataset-level variability. In contrast, the other methods showed narrower performance ranges but generally lower performance (median AUROC or ACC < 0.6). To reduce the influence of dataset-specific characteristics, we compared models after centering AUROC within each dataset and sampling setting. Among the evaluated methods, scDEAL showed the highest relative performance, with a median dataset-centered AUROC of 0.164 and a median normalized rank of 0.125, followed by DREEP with corresponding values of 0.038 and 0.333. Differences among methods therefore remained evident under dataset-matched comparisons (**Figure S3A and B**).

**Figure 2.**
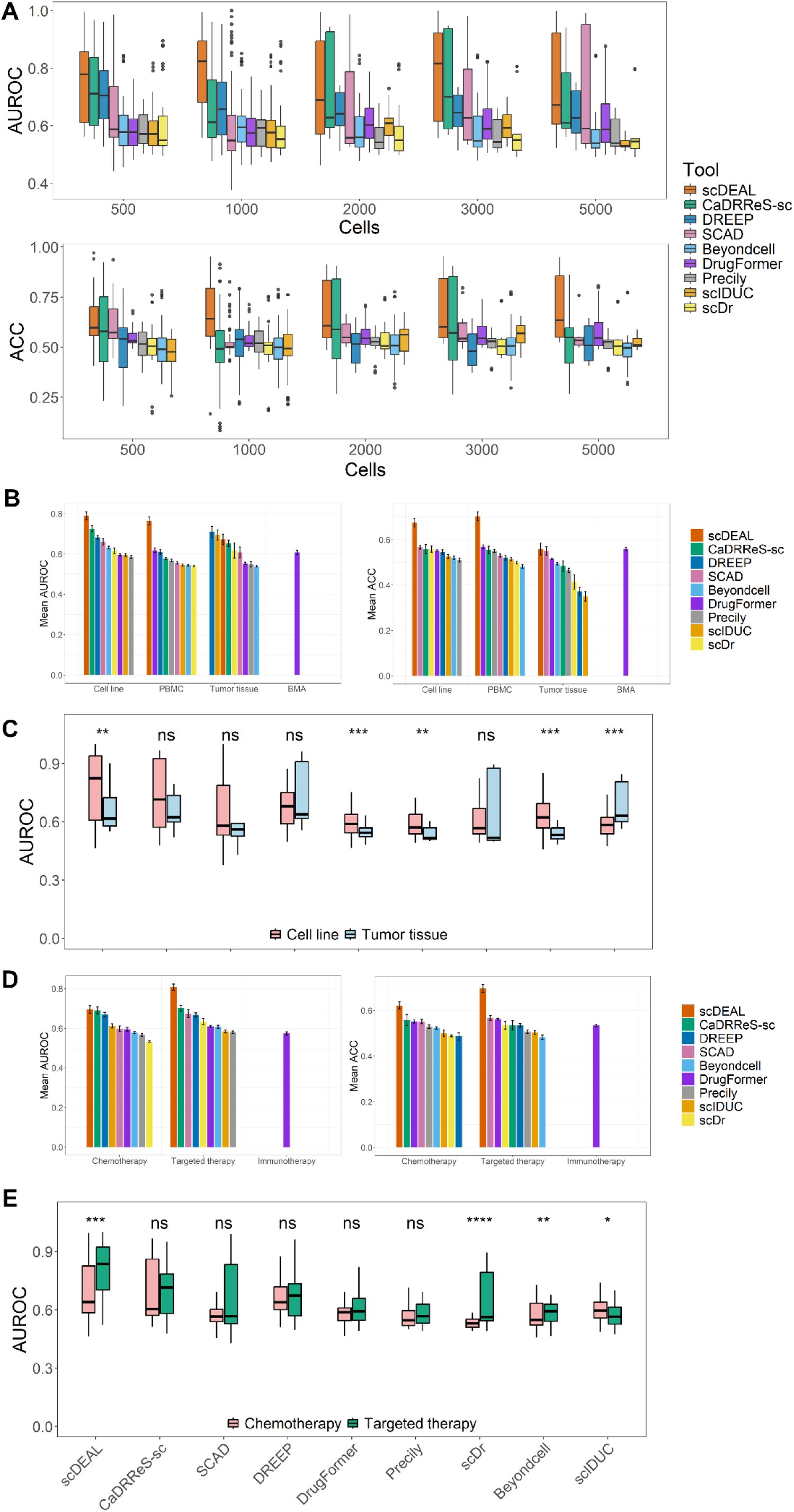
Performance on balanced datasets. **(A)** AUROC and ACC with different total numbers of cells in the datasets. **(B)** Mean AUROC (left) and accuracy (right) across datasets grouped by sample source. **(C)** Model performance comparisons grouped by sample source. (**D**) Mean AUROC (left) and accuracy (right) grouped by therapy type. (**E**) Model performance comparisons grouped by therapy type.

Further analysis of detailed evaluation metrics revealed that the performance of scDEAL was sensitive to sample size on specific datasets (**Figure S4**). For instance, in the GSE149124 dataset, scDEAL’s mean F1 jumped sharply from 0.33 at 500 cells to 0.73 at 2000 cells, indicating potential instability of its performance. We performed a Coefficient of Variation (CV) analysis based on three balanced subsampling under specific cell numbers (**Figure S5**). The results confirmed that, for the same dataset, scDEAL showed significantly higher CV values across multiple metrics than most other methods. For example, at 500 cells, scDEAL’s mean CVs for ACC and Precision were both ∼0.14. In contrast, other methods such as SCAD and DrugFormer exhibited greater stability, with their CVs generally ranging from 0.02 to 0.06(Wilcoxon rank-sum test, p < 0.05). This suggests that although scDEAL can achieve high predictive performance, it is highly sensitive to minor sampling variations, which may pose challenges to its robustness and reliability in practical applications.

To further explore how sample size affects model performance, we analyzed dataset sparsity, a key characteristic of single-cell data. The results showed that as the number of sampled cells increased, the overall sparsity of the datasets exhibited a clear upward trend (**Figure S6A**). This increased data sparsity significantly affects the performance of five of the nine evaluated algorithms (including scDEAL, Beyondcell, SCAD, and CaDRReS-sc) (**Figure S6B**). For the ACC metric, most methods showed patterns consistent with AUROC(**Figure S6C**). These results suggest that the effect of sample size on model performance may be mediated by dataset sparsity: as cell number increases, higher sparsity introduces substantial missing data that limits predictive performance.

We then stratified prediction performance by sample origin and drug type. Overall, model performance was significantly higher on cell-line datasets than on tumor tissues (**Figure S1C**; Wilcoxon rank-sum test, p = 2.8 × 10^-11^). scDEAL achieved mean AUROC values of 0.790 and 0.764 on cell-line and PBMC datasets, respectively, compared with 0.725 and 0.618 for the next-best method, whereas its mean AUROC decreased to 0.672 on tumor tissues, where performance across models ranged from 0.539 to 0.711 (**Figure 2B**). This difference may reflect that most methods were trained using cell-line-derived resources such as GDSC and CCLE, together with the greater cellular heterogeneity of tumor tissues [40]. At the model level, scDEAL, DrugFormer, Precily, and Beyondcell performed better on cell lines, whereas scIDUC showed the opposite trend (**Figure 2C**).

Performance was also significantly higher for targeted therapies than for chemotherapies (**Figure S1D**; Wilcoxon rank-sum test, p = 3 × 10^-7^). scDEAL performed similarly to other methods for chemotherapies but showed higher performance for targeted therapies (p < 0.01), with scDEAL, scDr, and Beyondcell generally performing better on targeted therapies and scIDUC showing the opposite pattern (**Figure 2D and E**). This may reflect the more specific molecular mechanisms of targeted therapies, whereas chemotherapeutic agents act through broader cellular processes that are less directly captured by gene-expression-based predictors [42, 43]. DrugFormer was the only evaluated method applicable to the immunotherapy datasets. Unlike most methods that depend on drug–cell associations derived from resources such as GDSC or CCLE, DrugFormer incorporates a knowledge graph based on biological relationships, including haploinsufficiency, which may enable it to represent gene-function relationships beyond conventional small-molecule response datasets [26].

To assess whether recent single-cell foundation models could improve prediction performance in the balanced setting, we compared scGPT and scFoundation with scDEAL under both within-dataset and cross-dataset settings. In within-dataset evaluation, scFoundation showed the strongest overall performance, with median AUROC, ACC, F1-score, precision, and recall of 0.982, 0.936, 0.935, 0.941, and 0.938, respectively, while the corresponding values for scGPT were 0.910, 0.825, 0.819, 0.882, and 0.764 (**Figure S7A**). Both foundation models generally exceeded scDEAL in this setting, with median Δ_AUROC_ values of 0.189 for scFoundation and 0.126 for scGPT (**Figure S7B**). Performance was lower under cross-dataset evaluation, with median AUROC values of 0.346 for scFoundation and 0.508 for scGPT, and corresponding median ΔAUROC values relative to scDEAL of −0.402 and −0.355. These results indicate that foundation models can achieve strong predictive performance in within-dataset settings, whereas their advantage is reduced under cross-dataset evaluation.

We next assessed the effects of sequencing platform and computational cost. AUROC values did not differ significantly between datasets generated using 10X Genomics and other platforms for most methods (Wilcoxon rank-sum test, p > 0.05; **Figure S8**). Runtime varied substantially across methods (**Figure S9**). Among Python-based methods, DrugFormer was the slowest, requiring 23.05 ± 0.54 min for 5,000 cells (p < 0.05), whereas Precily was the most time-consuming R-based method, requiring 118.67 ± 88.99 min and showing up to four-fold runtime differences across cancer types. Overall, R-based methods were slower than Python-based methods (p = 0.026), whereas memory usage did not differ. After excluding Precily, neither runtime nor memory usage differed between the two programming language groups (p = 0.99).

Taken together, increasing the cell number did not consistently improve model performance. Data sparsity rises with sample size, which in turn reduces prediction performance for most methods. The evaluated models are therefore highly context-dependent: methods such as scDEAL perform well in simpler settings (cell lines, targeted therapies) but lose their advantage in more complex contexts involving heterogeneous tumors and chemotherapies. Besides, foundation models such as scGPT and scFoundation performed strongly in within-dataset evaluation but showed reduced performance relative to scDEAL in cross-dataset evaluation, suggesting limited generalizability across datasets.

### Performance on Imbalanced Datasets

We next focused on scenarios that better reflect real cases, where resistant and sensitive cells often exhibit highly skewed proportions. Given that the original label distributions vary across datasets, we constructed imbalanced datasets by determining the maximal sample size for each target ratio, constrained by the available minority or majority cell counts and a global upper limit of 10,000 cells (**Figure 3A**).

**Figure 3.**
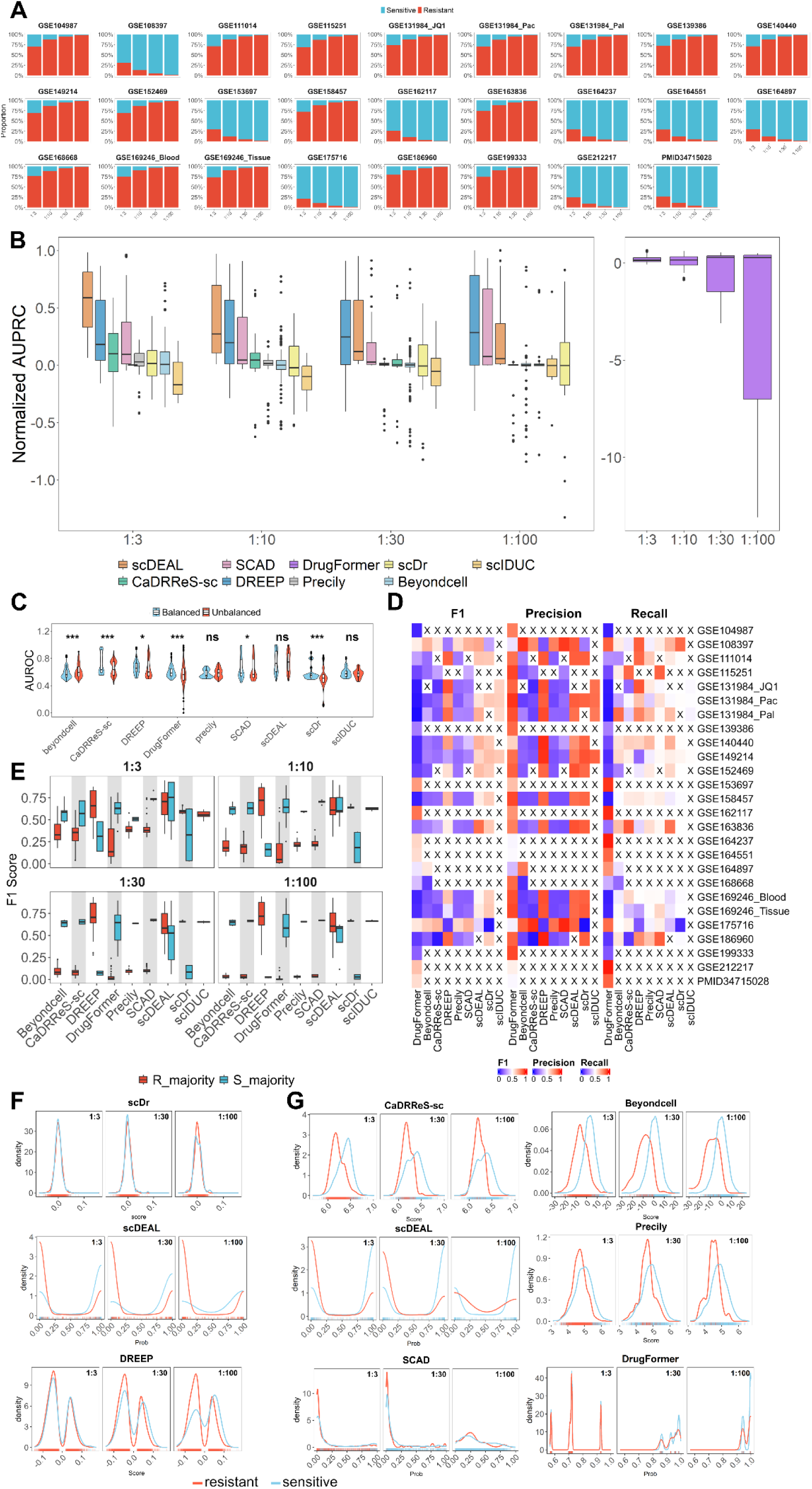
Performance on imbalanced datasets. **(A)** Proportions of resistant and sensitive cells across all datasets after imbalanced sampling at different majority/minority ratios. **(B)** Normalized AUPRC across different imbalanced ratios. **(C)** Comparison of AUROC between balanced and imbalanced scenarios. **(D** Mean F1-score, precision, and recall across all the imbalanced ratios. Crosses (×) indicate cases where a method failed to generate predictions. **(E)** F1 scores under resistant-majority versus sensitive-majority datasets across all imbalance ratios. **(F, G)** Density estimates of predicted scores for resistant and sensitive cells in dataset GSE163836 **(F)** and dataset GSE108397 **(G)**.

For most methods, AUROC remained relatively stable as class imbalance increased (**Figure S10A**), whereas only DrugFormer and scDEAL showed slight performance degradation under extreme imbalance. scDEAL consistently achieved the highest AUROC across all the imbalance ratios. In terms of ACC, DREEP achieved the highest performance across most ratios, followed closely by scDEAL (**Figure S10B**). However, when the imbalance became extreme (1:100), the ACC of scDEAL dropped markedly. In contrast, the ACC of other methods, which was already relatively low (0.5–0.6), did not show substantial fluctuations.

To enable fair comparisons across datasets with varying class imbalance ratios, we further introduced normalized AUPRC [44], which revealed substantial changes in the relative performance of the methods as class imbalance increased (**Figure 3B**, **Figure S11**). At moderate imbalance ratios of 1:3 and 1:10, scDEAL reached the highest median normAUPRC (0.587 and 0.271, respectively), compared with 0.182 and 0.196 for the next best method. However, at the ratio of 1:30, scDEAL’s performance declined, dropping to a median normAUPRC of 0.119, behind both DREEP (0.245) and DrugFormer (0.299). This downward trend continued at the extreme 1:100 ratio, where scDEAL’s median normAUPRC dropped to near-zero (0.056), as compared to DREEP (0.283) and DrugFormer (0.275) (**Figure 3B**). DrugFormer achieved moderate performance at the 1:3 and 1:10 ratios, but showed a sharp decline at the extreme 1:100 ratio. Notably, its predictions became inversely correlated with the true labels, suggesting a limited capacity to capture minority-class features under severe class imbalance. Similarly, we also compared models after centering normalized AUPRC within each dataset and imbalance ratio. scDEAL showed the highest relative performance, with a median dataset-centered normalized AUPRC of 0.194 and a median normalized rank of 0.143. DREEP also showed relatively strong performance, with corresponding values of 0.177 and 0.143 (**Figure S3C and D**).

We further assessed the overall stability of each method across balanced and imbalanced settings. **Figure 3C** showed that scDEAL consistently ranked at the top in both scenarios, achieving mean AUROC of 0.763 (balanced) and 0.743 (imbalanced). This minimal variation (Δ*_Auroc_*= 0.02; Wilcoxon rank-sum test, *p* > 0.05) highlights its limited sensitivity to class imbalance. Similarly, Precily and scIDUC demonstrated comparable stability, showing no significant performance shifts between the two settings (Δ*_Auroc_*= 0.013 and Δ*_Auroc_*= 0.019, respectively; Wilcoxon rank-sum test, *p* > 0.05). Despite this robustness, their overall performance was limited, achieving mean AUROCs of only 0.575/0.588 (Precily) and 0.594/0.574 (scIDUC) for balanced and imbalanced settings, respectively. In contrast, CaDRReS-sc, DREEP, DrugFormer, and scDr performed significantly better under balanced conditions than under imbalanced ones, whereas Beyondcell showed slightly improved performance in the imbalanced setting. To provide a more comprehensive evaluation beyond global ranking metrics, we further summarized F1-score, precision, and recall across all imbalanced datasets (**Figure 3D and Figure S12**). scDEAL showed relatively strong overall performance across these metrics, achieving a mean F1-score of 0.629 (mean precision: 0.875; mean recall: 0.568). scIDUC and DREEP followed with moderate performance, showing mean F1-scores of 0.622 and 0.609, respectively. In contrast, DrugFormer showed a distinct performance pattern: while it maintained a relatively high mean precision of 0.754, its mean recall was only 0.381, resulting in a notably low mean F1-score of 0.284. This disparity indicates that DrugFormer captures only a limited subset of high-confidence minority samples, consistent with its lower normAUPRC under severe class imbalance (**Figure 3B**).

We also evaluated scGPT and scFoundation using the same imbalanced datasets to explore whether foundation models could improve prediction under class imbalance. In the within-dataset setting, scFoundation achieved a median normalized AUPRC of 0.919 and ACC of 0.967, compared with 0.600 and 0.950 for scGPT, respectively (**Figure S7C**). Performance was lower in the cross-dataset setting, with median normalized AUPRC values of −0.017 for scFoundation and −0.010 for scGPT. At the dataset level, both models generally outperformed scDEAL in the within-dataset setting, with median Δ_normAUPRC_ values of 0.406 and 0.294 for scFoundation and 0.297 and 0.010 for scGPT at 1:3 and 1:30, respectively. In contrast, the corresponding values were negative under cross-dataset evaluation, reaching −0.605 and −0.095 for scFoundation and −0.638 and −0.081 for scGPT (**Figure S7D**). Similar to the balanced setting, foundation models generally outperformed scDEAL in within-dataset evaluation, although their advantage under class imbalance varied with imbalance severity. In contrast, both models performed worse than scDEAL in the cross-dataset setting, indicating that their strong within-dataset performance did not translate into improved cross-dataset generalization.

We further compared F1-scores between datasets dominated by resistant cells (R-majority) and sensitivity cells (S-majority) (**Figure 3E**). Under moderate imbalance (1:3), although performance differences between the two dataset types were observed for some methods, the overall disparities were relatively small (average Δ*_F_*_1_= 0.234). However, under severe imbalance such as 1:100, pronounced divergence emerged for most methods. For example, CaDRReS-sc, SCAD, Precily, DrugFormer, and Beyondcell achieved substantially higher F1-scores on S_majority datasets than on R_majority datasets (average Δ*_F_*_1_= 0.663, 0.634, 0.626, 0.618, and 0.613, respectively). Conversely, DREEP and scDr performed better on R-majority datasets (average Δ*_F_*_1_= 0.683 and 0.637). In contrast, the performance of scDEAL remained relatively stable across both dataset types even under extreme imbalance (average Δ*_F_*_1_= 0.157), indicating less sensitivity to the direction of class imbalance than most other methods.

Finally, to further examine prediction-score distributions at the dataset level, we selected GSE163836 (R-majority) and GSE108397 (S-majority) as representative datasets, as the drugs tested in these datasets are supported by most evaluated tools. On GSE163836, although scDr achieved the competitive performance (average F1 = 0.642), its predicted drug response scores did not clearly separate resistant and sensitive cells, as reflected by highly overlapping density distributions (**Figure 3F**). This is likely because its prediction scores are conservative, being compressed into a narrow range, which may account for its apparently stable performance under class imbalance setting. DREEP showed similar behavior to scDr, with limited separability but slightly improved discrimination. In contrast, scDEAL demonstrated clearer discrimination. Under moderate imbalance, resistant cells were mainly predicted with low-probabilities, whereas sensitive cells were predicted with high-probabilities.

Nevertheless, at the extreme 1:100 ratio, resistant cells slightly overlapped with sensitive cells in the high-probability region, suggesting an accumulation of false positives driven by the overwhelming size of the majority class. Consequently, when the ratio shifted from 1:3 to 1:100, the precision decreased from 0.969 to 0.871. Although the recall remained relatively stable (declining from 0.857 to 0.813), the increased number of false positives ultimately reduced the overall F1-score from 0.898 to 0.791(**Figure S12**).

On GSE108397, scDEAL again showed strong discrimination, although under the 1:100 setting, the density of sensitive cells partially shifted toward the low-probability region (**Figure 3G**). CaDRReS-sc and Beyondcell also showed a modest separation between resistant and sensitive cells. Precily demonstrated observable class separation across varying class ratios (average AUROC = 0.656). However, the observed scoring pattern was opposite to the intended interpretation of the model output, where higher scores are expected to indicate stronger resistance. In our evaluation, sensitive cells received slightly higher scores (median = 4.894) than resistant cells (median = 4.673). In contrast, SCAD and DrugFormer showed limited discrimination between resistant and sensitive cells.

In summary, model performance under class imbalance depended on both the severity and direction of class imbalance. scDEAL maintained the highest AUROC and relatively strong F1 performance under moderate imbalance settings. Under severe imbalance, however, methods such as DREEP and SCAD outperformed scDEAL. Foundation models achieved improved performance in within-dataset evaluations but showed performance reductions in the cross-dataset setting. Most methods also showed substantial differences between R-majority and S-majority datasets, whereas scDEAL was less sensitive to the direction of class imbalance. These results indicate that no single method consistently dominated across all metrics and imbalance levels, and the overall performance deterioration under severe imbalance highlights the limited ability of current models to identify rare resistant or sensitive cell populations.

### Performance on Lineage Tracing Datasets

To further assess model performance in a clinically relevant setting, we conducted benchmarking on two independent lineage-tracing datasets, ReSisTrace [20] and Watermelon [21]. In both datasets, lineage information enables pre-treatment cells to be classified as pre-sensitive or pre-resistant according to the subsequent fate of their corresponding lineages, providing experimentally supported response labels that are not available in conventional scRNA-seq datasets.

We first evaluated the models using only pre-treatment transcriptomic profiles in the ReSisTrace dataset. In this setting, lineage-tracing labels allow pre-treatment cells to be classified as pre-sensitive or pre-resistant according to the fate of their matched sister cells after treatment. Model performance was generally limited, with mean AUROC values ranging from 0.502 to 0.622 across the evaluated methods. Among them, scDEAL achieved the highest mean AUROC of 0.622, with an ACC of 0.588 and an F1-score of 0.593 (**Figure 4A**). UMAP visualizations further showed extensive overlap between pre-sensitive and pre-resistant cells, and predictions from most methods were highly intermixed across the UMAP projection (**Figure 4B**). For comparison, when the same dataset was evaluated under the conventional pre-treatment versus post-treatment setting, performance was substantially higher. scDEAL achieved a mean AUROC of 0.869, followed by Beyondcell at 0.779, while CaDRReS-sc, SCAD, and Precily reached 0.699, 0.698, and 0.656, respectively (**Figure S13A**). Predicted scores also showed partial separation between the pre- and post-treatment populations (**Figure S13B**).

**Figure 4.**
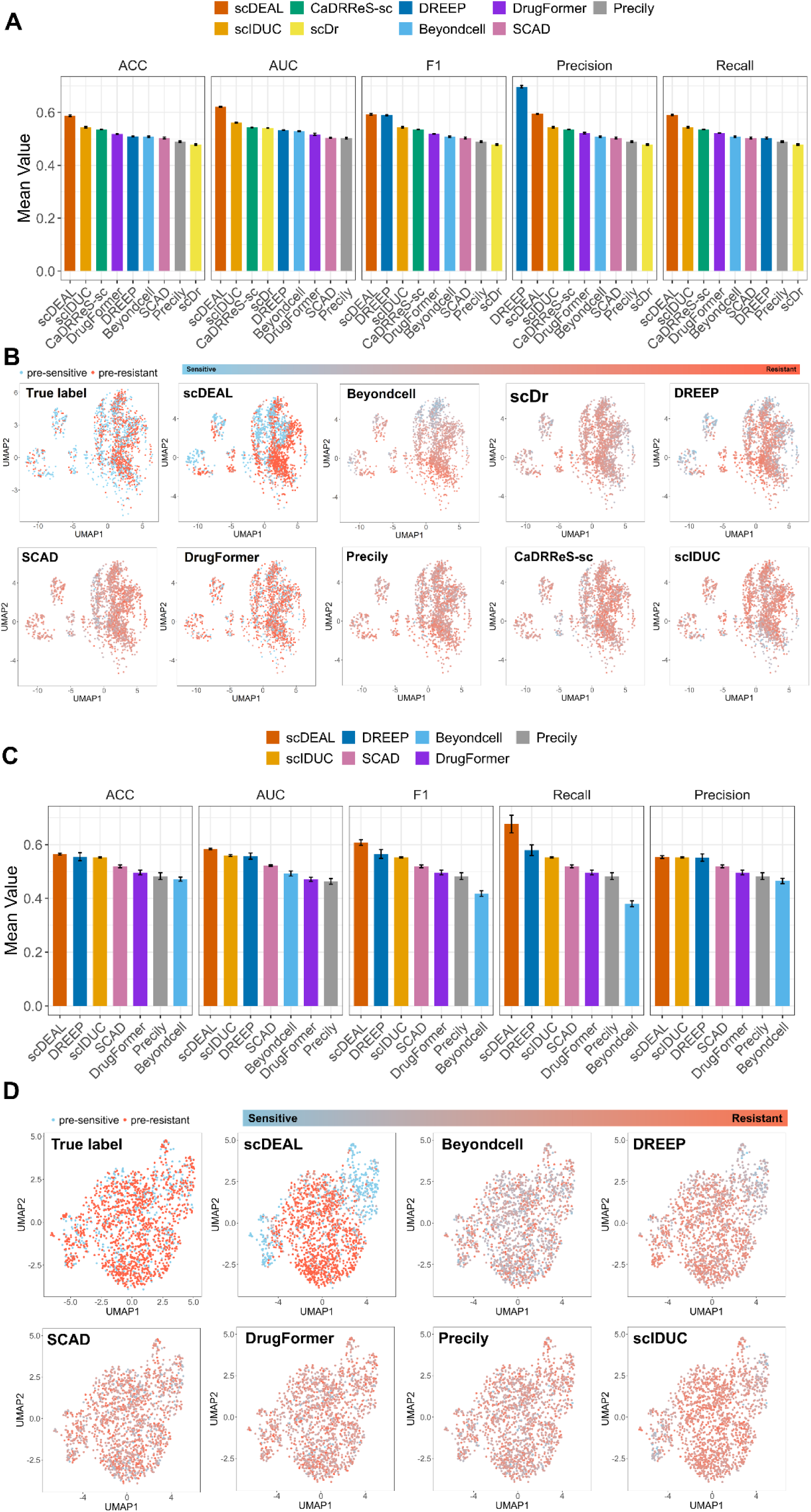
Performance on lineage-tracing single-cell drug response data. **(A, B)** Results on the ReSisTrace dataset for distinguishing pre-sensitive and pre-resistant cell populations using pre-treatment data only. **(A)** Mean accuracy, AUROC, F1 score, precision, and recall across all evaluated methods. **(B)** UMAP visualization of ground truth labels and model-predicted scores. **(C, D)** Corresponding results on the Watermelon dataset. **(C)** Mean accuracy, AUROC, F1 score, precision, and recall across all evaluated methods. **(D)** UMAP visualization of ground truth labels and model-predicted scores.

A similar pattern was observed in the Watermelon dataset. When only D0 pre-treatment cells were used to distinguish pre-sensitive from pre-resistant states, mean AUROC values across the evaluated methods ranged from 0.463 to 0.584, with scDEAL showing the highest mean AUROC of 0.584, together with an ACC of 0.564 and an F1-score of 0.608 (**Figure 4C**). Model-predicted scores again showed substantial overlap between pre-sensitive and pre-resistant cells (**Figure 4D**). In contrast, prediction performance increased when pre-treatment cells were compared with post-treatment resistant cells. scDEAL achieved a mean AUROC of 0.997, followed by DREEP at 0.943, compared with a highest mean AUROC of only 0.584 in the intrinsic pre-sensitive versus pre-resistant task (**Figure S13C and D**). Across both lineage-tracing datasets, treatment-associated states were therefore more readily distinguished than lineage-defined pre-existing resistance states.

We also examined whether recent single-cell foundation models could improve prediction of pre-existing resistance. On ReSisTrace, scFoundation and scGPT achieved mean AUROC values of 0.662 and 0.602, respectively, compared with 0.622 for scDEAL. On Watermelon, the mean AUROC was 0.598 for scFoundation, 0.567 for scGPT, and 0.584 for scDEAL (**Figure S7E**). Although scFoundation achieved the best performance on both datasets, its discrimination remained modest. UMAP visualization of predicted scores similarly showed only partial correspondence with the lineage-derived response labels (**Figure S7F and G**). Thus, the limited predictability of pre-existing resistance was also observed with foundation-model-based representations.

In ReSisTrace, we additionally examined whether model-predicted labels recapitulated the underlying transcriptional differences between response groups. Under the conventional pre-versus post-treatment setting, scDEAL correctly classified approximately 75% of cells and showed a strong positive correlation between predicted and ground-truth differential-expression patterns (**Figure S14A and D**, Pearson R = 0.78). In contrast, Beyondcell and CaDRReS-sc exhibited pronounced label inversion and negative correlations with the ground truth (R = −0.68 and −0.61, respectively), despite Beyondcell achieving the highest DEG-overlap Jaccard index of 0.53 (**Figure S14B**). Agreement was weaker in the intrinsic pre-sensitive versus pre-resistant task (**Figure S15A**). CaDRReS-sc achieved the highest DEG-overlap Jaccard index of only 0.087, followed by scDEAL at 0.051, while their fold-change correlations with the ground truth were R = 0.50 and approximately 0, respectively (**Figure S15B and D**). These results further reflect the difficulty of recovering pre-existing resistance-associated transcriptional signals from baseline profiles.

To investigate whether the limited predictive performance could reflect weak transcriptional differences already present before treatment, we directly characterized pre-sensitive and pre-resistant cells in both lineage-tracing datasets. In ReSisTrace, 10 DEGs were identified between the two populations (**Figure S16A**), with enrichment of ribosome, MYC-target, proteasome, oxidative-phosphorylation, and cell-cycle-related programs in pre-resistant cells, whereas pre-sensitive cells were associated with TNFα/NF-κB signaling, apoptosis, and interferon-γ response (**Figure S16B**) [45–47]. In Watermelon dataset, 64 DEGs were identified between pre-sensitive and pre-resistant cells, with enrichment of E2F targets, homologous recombination, G2M checkpoint, MYC targets, oxidative phosphorylation, and mTORC1 signaling in pre-resistant cells, while pre-sensitive cells showed enrichment of pathways including ribosome, xenobiotic metabolism, glutathione metabolism, lysosome, and hypoxia (**Figure S17B, C and D**) [48–50]. In both datasets, these intrinsic differences were substantially weaker than treatment-associated changes, as reflected by the lower DEG numbers and smaller absolute log2 fold changes in the pre-sensitive versus pre-resistant contrasts (**Figure S16C and Figure S17E**).

Taken together, these results show that predicting pre-existing resistance from untreated baseline transcriptomes is more difficult than distinguishing treatment-associated transcriptional states. The inclusion of scGPT and scFoundation did not notably change this overall pattern. Transcriptomic analyses further showed that pre-sensitive and pre-resistant cells exhibit detectable but relatively subtle differences compared with the stronger changes associated with treatment. The limited strength of these pre-existing transcriptional signals may therefore partly explain the difficulty of identifying resistant cells before treatment using current transcriptome-based drug-response prediction models.

### Architecture-dependent training-label effects, component contributions and biological interpretation of scDEAL performance

To examine the factors underlying the performance of scDEAL, we first compared the training labels used by scDEAL and SCAD. Across 34 drugs with at least 30 common cell lines, the two labeling systems showed substantial overall concordance, although the degree of agreement varied across drugs (**Figure 5A and Figure S18A**).

**Figure 5.**
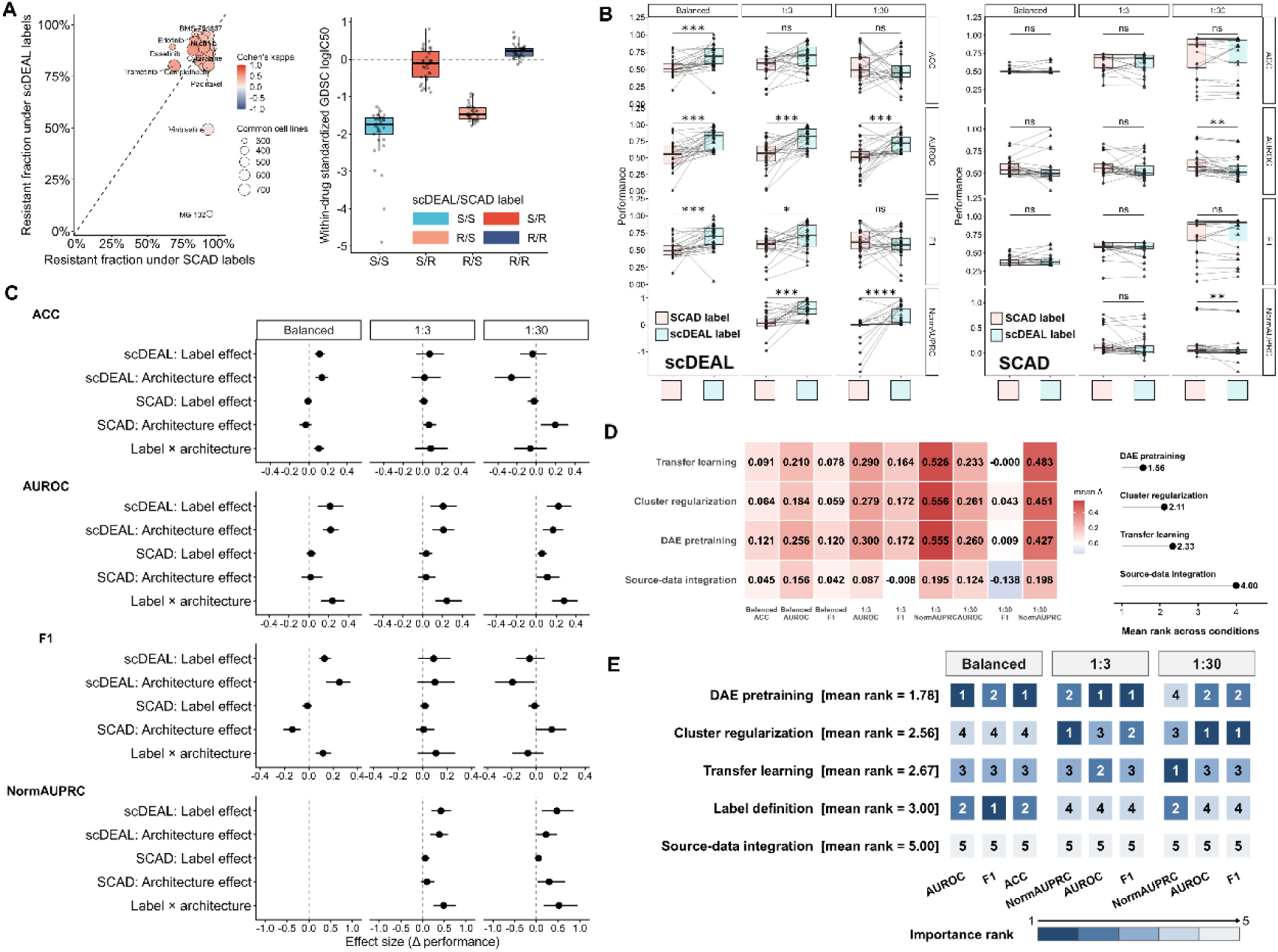
Dissecting the contributions of model components to scDEAL performance. **(A)** Comparison of drug-response labels defined by SCAD and scDEAL. **(B)** Performance of scDEAL and SCAD using their native versus substituted training labels under balanced and imbalanced settings. **(C)** Label, architecture, and label–architecture interaction effects across different evaluation settings. **(D)** Contribution of individual architectural components to scDEAL performance. (**E**) Overall contribution ranking of scDEAL model components across evaluation settings. ns, not significant; * P < 0.05; ** P < 0.01; *** P < 0.001; **** P < 0.0001.

Concordant labels (S/S and R/R) accounted for the majority of common cell lines for 33/34 drugs, and Cohen’s kappa was positive for 30/34 drugs, with a median κ of 0.22. However, several drugs showed marked differences in label assignment. For example, approximately 90% of the common cell lines were labeled as resistant to MG-132 by SCAD, compared with ∼8% by scDEAL, while the corresponding resistant fractions for Vinblastine were ∼90% and ∼50%, respectively. The four label combinations also showed distinct GDSC logIC50 distributions: the median within-drug standardized logIC50 values were −1.740, −0.111, −1.470, and 0.226 for S/S, S/R, R/S, and R/R, respectively (**Figure 5A**).

We next replaced the training labels between scDEAL and SCAD and evaluated their performance under balanced, 1:3, and 1:30 settings (**Figure 5B and Figure S18B**). Label substitution significantly affected scDEAL in 8 of the 11 evaluated metric– setting combinations, whereas only 2 of 11 comparisons were significant for SCAD. Specifically, under scDEAL, the native scDEAL labels produced higher AUROC in all three settings (all P < 0.001), and higher normalized AUPRC at both 1:3 (P < 0.001) and 1:30 (P < 0.0001). Significant improvements were also observed for ACC in the balanced setting (P < 0.001) and for F1 in the balanced and 1:3 settings (P < 0.001 and P < 0.05, respectively). In contrast, replacing SCAD labels with scDEAL labels produced no significant changes in the balanced or 1:3 settings. At 1:30, the native SCAD labels gave higher AUROC and normalized AUPRC than the substituted labels (both P < 0.01).

Effect decomposition further showed that the influence of training labels depended on the model architecture (**Figure 5C**). The scDEAL-specific label effect on AUROC remained positive across all three settings, with effect sizes of approximately 0.21– 0.22, whereas the corresponding SCAD label effects remained close to zero. For normalized AUPRC, the scDEAL label effect increased to approximately 0.4–0.5 under the imbalanced settings, while the SCAD label effect remained near zero.

Architecture effects were more dependent on the evaluation setting. For example, the mean scDEAL architecture effect on ACC changed from approximately 0.15 in the balanced setting to approximately −0.25 at 1:30, with a similar change observed for F1. In contrast, the label × architecture interaction remained positive for AUROC (∼0.25 across settings) and normalized AUPRC (∼0.45–0.55), indicating that the effect of the training-label definition was strongly dependent on the architecture.

We then performed ablation analysis to quantify the contribution of individual scDEAL components (**Figure 5D**). DAE pretraining showed the largest overall contribution among the four architectural components, with a mean rank of 1.56, followed by cluster regularization (2.11), transfer learning (2.33), and source-data integration (4.00). DAE pretraining showed consistent contributions across settings, with mean ΔAUROC values of 0.256, 0.300, and 0.260 in the balanced, 1:3, and 1:30 settings, respectively. Under class imbalance, cluster regularization and transfer learning showed particularly large effects on normalized AUPRC, with Δ values of 0.556 and 0.526 at 1:3, and 0.451 and 0.483 at 1:30, respectively. In contrast, source-data integration showed the smallest overall contribution, with a mean rank of 4.00 and relatively small performance changes across most conditions. Its effect on F1 was slightly negative at 1:3 (Δ = −0.008) and further decreased at 1:30 (Δ = −0.138).

When training-label definition was further included in the overall component ranking, DAE pretraining remained the highest-ranked component with a mean rank of 1.78, followed by cluster regularization (2.56), transfer learning (2.67), and label definition (3.00), whereas source-data integration ranked last (5.00) (Figure 5E). DAE pretraining ranked within the top two components in 8 of the 9 metric–setting combinations and ranked first in four. In comparison, label definition ranked first in one condition and within the top two in four of nine conditions. Source-data integration ranked fifth in all nine conditions. Together, these analyses indicate that scDEAL performance is shaped by both its training-label definition and architectural design, with DAE pretraining, cluster regularization, and transfer learning providing the largest overall contributions.

Having identified the technical factors contributing to scDEAL performance, we next examined whether its predictions were also associated with biologically interpretable drug-related cellular states in an independent pancreatic ductal adenocarcinoma (PDAC) organoid dataset. The dataset comprised DMSO controls and cells treated with paclitaxel or trametinib. Drug treatment induced substantial transcriptomic remodeling, with clear separation between the control and drug-treated conditions in unsupervised dimensionality reduction analyses (**Figure6A–C**). In total, 7,301 cells from two independent samples were analyzed, with 1,647–2,948 cells per condition (**Figure 6D**).

**Figure 6.**
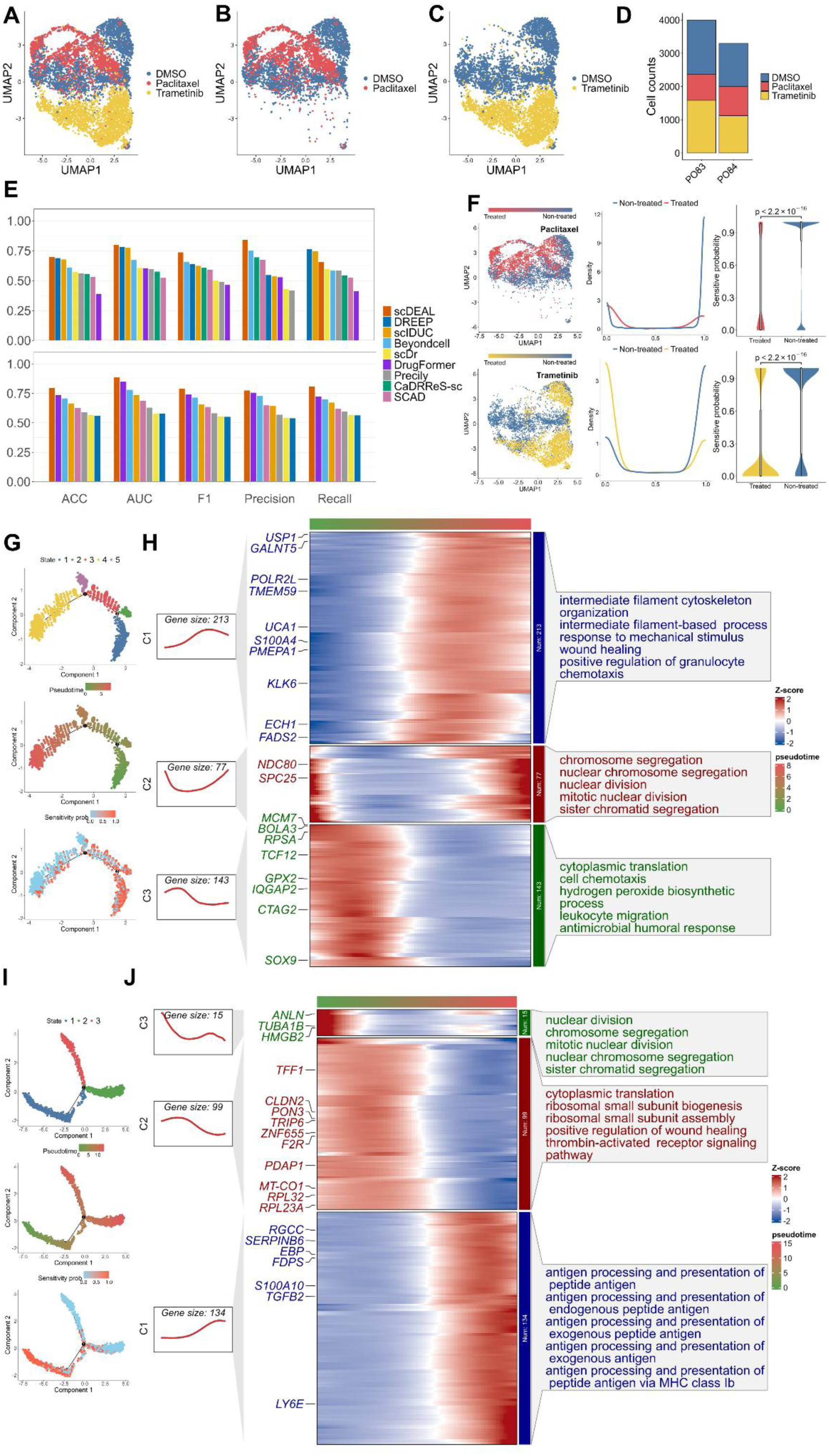
Independent validation of scDEAL in a PDAC cohort reveals drug-induced transcriptional state transitions. **(A–C)** UMAP visualization of the PDAC dataset colored by treatment conditions: **(A),** paclitaxel vs. DMSO **(B),** and trametinib vs. DMSO **(C). (D)** Cell counts across treatment groups and samples. **(E)** Performance metrics of the benchmarked methods under paclitaxel (top) and trametinib (bottom) treatments. **(F)** Predicted drug response scores shown by UMAP (left), density (middle), and violin plots (right). **(G-H)** Pseudotime trajectories **(G)** and dynamic gene expression with functional enrichment **(H)** for the paclitaxel-treated condition. In (**G**), cells are colored by cell state (top), pseudotime (middle), and scDEAL-predicted sensitivity probability (bottom; 1 = sensitive, 0 = resistant). **(I–J)** Pseudotime trajectories (**I**) and dynamic gene expression with functional enrichment **(J)** for the trametinib-treated condition. In (**I**), cells along the trajectory are colored by cell state (top), pseudotime (middle), and predicted sensitivity probability by scDEAL (bottom; 1 = sensitive, 0 = resistant).

scDEAL also showed relatively strong predictive performance in this dataset, achieving an accuracy of 0.698, AUROC of 0.801, and F1-score of 0.736 for paclitaxel, and corresponding values of 0.776, 0.834, and 0.771 for trametinib (**Figure 6E**). Predicted response scores differed markedly between DMSO and drug-treated cells for both treatments (Wilcoxon rank-sum test, P < 2.2 × 10⁻¹⁶; **Figure 6F**).

Consistent with these results, visualization of the prediction landscapes and confusion matrices showed clearer separation and more balanced classification for scDEAL than for several other methods (**Figures S19 and S20**). These observations indicate that scDEAL predictions reflected biologically meaningful treatment-associated cellular variation in this independent dataset.

To further characterize the biological processes associated with the scDEAL predictions, we performed pseudotime trajectory analysis. In the paclitaxel condition, cells with higher predicted sensitivity were enriched near the beginning of the inferred trajectory, whereas cells with lower predicted sensitivity were more frequently observed toward terminal states (**Figure 6G, top**). A comparable progression was observed in the trametinib condition **(Figure 6I, top**). Consistently, predicted response probabilities changed gradually along the inferred trajectories for both treatments (**Figure 6G and I, middle and bottom**), suggesting that the model scores tracked continuous treatment-associated transcriptional changes.

Pseudotime-associated genes further revealed treatment-specific biological programs. Under paclitaxel treatment, late-stage genes were enriched in intermediate filament cytoskeleton organization and responses to mechanical stimuli, while another gene cluster showed dynamic changes in chromosome segregation and nuclear division, consistent with cell-cycle perturbation associated with paclitaxel treatment (**Figure 6H**) [51, 52]. Genes involved in cytoplasmic translation and cell chemotaxis showed higher expression at earlier pseudotime stages. Under trametinib treatment, late-stage genes were enriched in antigen processing and presentation, whereas other dynamic clusters involved ribosome assembly, cytoplasmic translation, mitosis, and chromosome segregation (**Figure 6J**) [53, 54]. These patterns indicate that the predicted response gradients were associated with coherent transcriptional programs related to the biological effects of the corresponding treatments.

We finally stratified cells according to the median scDEAL prediction score and examined drug-associated pathways. For paclitaxel, known drug targets such as TUBB1, BCL2, and NR1I2 showed sparse expression and no clear differences between the two predicted groups (**Figure S21A**). In contrast, pathway-level analysis showed higher G2/M checkpoint, microtubule-stability, and paclitaxel-resistance signature scores among cells with lower predicted sensitivity, whereas apoptosis showed a modest increase among cells with higher predicted sensitivity (**Figure S21B**). Together, these analyses suggest that, beyond its dependence on training-label definition and architectural components, scDEAL predictions were associated with treatment-related transcriptional and pathway-level variation in an independent PDAC organoid dataset.

## Discussion

Despite the rapid development of computational methods for single-cell drug response prediction, the lack of comprehensive evaluation and high-quality ground-truth validation makes it difficult to fully assess their performance. In this study, we conducted a systematic benchmarking analysis of nine representative single-cell drug response prediction methods using large-scale public datasets and two independent lineage-tracing scRNA-seq datasets. We further evaluated two recent single-cell foundation models as complementary benchmarks. Notably, the lineage-tracing datasets provide experimentally supported response labels that enable evaluation of pre-treatment resistance prediction. By evaluating model performance across multiple scenarios, including balanced, imbalanced, and lineage-tracing datasets, we not only quantified differences among these methods but also examined how data heterogeneity, class imbalance, and the definition of the prediction task influence model performance.

Our results highlighted the impact of training data sources and therapeutic mechanisms on model performance. Increasing the number of sampled cells did not consistently improve prediction performance across methods. Instead, the accompanying increase in matrix sparsity was associated with negative changes in model performance, suggesting that larger single-cell datasets do not necessarily provide more usable predictive information. Although scDEAL showed competitive performance across several benchmark settings, its relatively high variability across datasets and subsampling rounds indicates that its predictions remain sensitive to dataset composition and sampling variation. These findings suggest that current single-cell drug response prediction models are still affected by data sparsity and dataset-specific characteristics. We also observed that most models perform better on cell-line datasets than on tumor tissue samples, and they generally achieve higher predictive accuracy for targeted therapies than for chemotherapeutic agents. This may partly reflect the fact that most existing prediction tools are constructed using bulk-level transcriptomic profiles from large-scale cell-line resources such as GDSC and CCLE. Although many methods have adopted transfer learning or domain-adaptation strategies to enable bulk-to-single-cell knowledge transfer [55], the performance decline observed in clinical samples indicates that these approaches may not fully capture the biological differences between in vitro training systems and *in vivo* tumors [4]. Clinical tumor tissues are characterized by intrinsic transcriptional heterogeneity and subpopulation complexity that commercial cell lines often cannot fully recapitulate. In addition, chemotherapeutic agents typically act through broad cytotoxic mechanisms involving diverse stress-response pathways, whereas targeted therapies operate through more specific molecular perturbations. This mechanistic distinction may contribute to less distinct transcriptional response patterns for chemotherapy than for targeted therapies. Also, predicting immunotherapy responses remains a major bottleneck. With the exception of DrugFormer, which demonstrated only limited predictive capability, most of the current models are not optimized for immunotherapy. One possible reason is that most models rely on *in vitro* cell-line training datasets that lack the information about tumor microenvironment and cell-cell interactions. Addressing this domain shift from *in vitro* to *in vivo* requires biologically informed transfer strategies that emphasize mechanism-aware and context-dependent feature learning. Recent studies such as PERCEPTION [56] have begun to explore such biologically informed transfer learning frameworks. Although not included in our benchmarking due to its limited drug coverage, this study highlights a future direction that moves from statistical alignment toward mechanism-aware modeling to better capture the complexity of tumor biology. Our evaluation of scGPT and scFoundation further showed that foundation-model representations can achieve strong performance within individual datasets, but their advantage was reduced under cross-dataset evaluation. This suggests that pretrained representations alone may not fully resolve the challenge of cross-dataset generalization.

The class imbalance commonly observed in real-world clinical data represents a major bottleneck that limits the clinical translation of prediction models [57]. In actual clinical settings, resistant cells can exist as rare subpopulations, and such highly skewed data distributions can significantly affect the performance of machine learning algorithms [58, 59]. Our results show that under extreme imbalance ratios such as 1:100, prediction performance decreased for many methods. scDEAL maintained relatively strong performance under moderate imbalance and showed less sensitivity to the direction of class imbalance than most other methods. However, this advantage was not consistent across all imbalance levels and metrics. Under severe imbalance, its normalized AUPRC declined, and DREEP and SCAD achieved higher values. Similarly, scGPT and scFoundation performed strongly in within-dataset evaluations but showed marked performance reductions under cross-dataset evaluation. These results indicate that no single method consistently dominated across different imbalance settings.

One of the key insights of our study is derived from the benchmarking using lineage-tracing data. Across both ReSisTrace and Watermelon, under the conventional evaluation setting, where pre-treatment cells are compared with post-treatment surviving cells, most models achieve relatively high accuracy, likely because drug perturbation induces substantial transcriptional changes that make the two samples easier to distinguish. However, such performance may partly reflect the models’ ability to detect treatment-associated transcriptional differences instead of directly indicating their ability to predict pre-existing resistance. A similar contrast between conventional and pre-treatment prediction was observed for scGPT and scFoundation, suggesting that this pattern was not restricted to the original task-specific models.

More importantly, our concordance analysis showed that high DEG overlap alone can be misleading, because models with inverted labels may still recover similar DEG sets but assign the opposite direction of transcriptional changes. Therefore, model evaluation should not rely only on classification performance or DEG overlap. The direction of transcriptional changes should also be considered when assessing the biological consistency of model predictions [60].

When the task is shifted to a more clinically relevant setting—predicting a cell’s eventual fate before drug treatment—performance for most models declined markedly and remained relatively modest in both lineage-tracing datasets. This difficulty may reflect both limitations of current models and the relatively weak transcriptional differences associated with future drug response before treatment. Our bioinformatics analyses showed that pre-sensitive and pre-resistant cells contained detectable differential expression and pathway-level signals in both ReSisTrace and Watermelon, but these differences were weaker than the transcriptional changes observed following treatment. These relatively subtle intrinsic signals may partly explain the limited predictive performance of current models when using pre-treatment transcriptomes. Recent evidences have also highlighted that identical clones can take on divergent transcriptional paths and exhibit distinct drug sensitivity or resistance phenotypes, regardless of whether they are exposed to drugs [61]. This intrinsic cellular plasticity makes it inherently challenging to accurately predict eventual treatment outcomes based solely on a snapshot of transcriptional signatures at a given time in the past, even among genetically identical sister cells. Besides, intrinsic resistance may also depend on molecular information that is not fully captured by pre-treatment transcriptomes. Genetic alterations and chromatin states may therefore provide complementary signals for predicting future drug response [62, 63]. Consequently, current representation learning approaches may have limited ability to identify the subtle and intrinsic transcriptional features associated with cell-fate decisions under treatment. Although scFoundation showed a modest improvement in some lineage-tracing evaluations, neither the foundation models nor the task-specific methods consistently achieved strong discrimination of pre-existing resistance across both datasets. These results highlight the importance of distinguishing adaptive resistance from intrinsic resistance at the single-cell level and highlight the value of lineage-tracing data for identifying transcriptional features primed to drug resistance.

Finally, our training-label substitution and ablation analysis provided further insight into the factors contributing to scDEAL performance. Although the two label definitions showed majority agreement for most drugs, the degree of agreement varied across compounds, and replacing the labels had a substantially larger effect under the scDEAL architecture than under SCAD. Effect decomposition further showed a positive interaction between label definition and model architecture. At the same time, the ablation analysis demonstrated substantial contributions from the architectural components of scDEAL. DAE pretraining showed the largest overall contribution, followed by cluster regularization and transfer learning, whereas the contribution of training-label definition was more dependent on the evaluation setting. These results suggest that both training-label definition and architectural design contribute to scDEAL performance. As a complementary analysis, our in-house PDAC organoid case study showed that scDEAL-predicted response states were associated with treatment-related cellular transitions and pathway changes, supporting the biological relevance of its predictions in a treatment-associated setting. These analyses indicate that scDEAL performance reflects contributions from training-label definition and architectural components, while its predictions retain biologically meaningful treatment-associated patterns.

Although our study provides a comprehensive benchmarking analysis, several limitations remain. First, although Watermelon provides an independent lineage-tracing system in addition to ReSisTrace, the evaluation of pre-existing resistance is still limited to two lineage-tracing datasets. Additional datasets covering different cancer types, therapeutic mechanisms, and biological contexts will be needed to determine how broadly the observed patterns generalize. Second, although we included scGPT and scFoundation in the revised benchmark, these represent only two single-cell foundation models and were evaluated using pretrained representations followed by a common downstream classifier. Other foundation models or alternative fine-tuning strategies may yield different results. Third, our current evaluation only focuses on single-drug interventions. However, combination therapy is the standard treatment strategy in clinical practice. Although some studies have started to predict cell responses to drug combinations at the single-cell level [24, 28, 64], a systematic benchmark of these methods is missing. Expanding the current evaluation to test model accuracy on drug combinations is highly necessary in the future.

## Acknowledgement

The author would like to thank the FIMM High-Throughput Biomedicine unit for access to drugs and drugging technology, the FIMM Single-Cell Analytics unit for single cell transcriptomic services, and the FIMM High-Content Image Analysis unit for help with automated confocal microscopy of organoids. All FIMM units are supported by HiLIFE and Biocenter Finland. The authors also thank the iCANDOC Precision Cancer Medicine (PCM) pilot for supporting this project. ChatGPT (OpenAI, USA) was used with careful consideration to improve the language and readability of the manuscript text.

This work was supported by The Sigrid Jusélius Foundation (J. Tang); The Cancer Foundation Finland (J. Tang); Research Council of Finland projects 317680, 320131, 351196 (J. Tang) and 351198 (H. Seppänen); ReSisTrace funding decisions 351165 (J. Tang); This study was co-funded by the European Union (J. Tang: European Research Council project DrugComb No. 716063 and EATRIS-CONNECT No. 101130349). Views and opinions expressed are however those of the author(s) only and do not necessarily reflect those of the European Union or the European Research Council. Neither the European Union nor the granting authority can be held responsible for them.

## Author contributions

L.S. and J.T. conceived and designed the study. L.S., X.S., and S.Z. performed the benchmarking analyses. A.H. and M.V.-K. performed the PDAC single-cell experiments and initial data preprocessing. L.S. and X.S. performed the bioinformatics analyses of the PDAC single-cell data. L.S. and J.T. wrote the initial draft of the manuscript. J.E., H.M., H.S., B.S., M.L., and J.T. reviewed and edited the manuscript. J.T. supervised the study. All authors read and approved the final manuscript.

## Data availability

The publicly available datasets used for benchmarking in this study are described in detail in the Materials and Methods section and are available through the DRMREF database (https://ccsm.uth.edu/DRMref/). The ReSisTrace lineage-tracing single-cell RNA-seq dataset used for ground-truth evaluation was generated in our previous study and is publicly available from the Gene Expression Omnibus under accession number GSE223003. The Watermelon lineage-tracing single-cell RNA-seq dataset is publicly available from the Gene Expression Omnibus under accession number GSE150949. The raw single-cell RNA-seq data generated from the PDAC organoids in this study will be deposited in a public repository and made fully accessible before publication. During the initial peer review process, these raw data are available from the corresponding author upon reasonable request. The processed data files required to reproduce the analyses are currently available to editors and reviewers through Google Drive (https://drive.google.com/file/d/1qUr4buFPajm3bRUvZabjPUtfzIhcyEvM/view?usp=sharing). These files, together with the benchmarking code, will be deposited in a public research repository and made publicly available before publication.

## Code availability

Figure-generation scripts are available at GitHub: https://github.com/daleeshen/scDrugPred_benchmark. The complete reproducibility package is available to reviewers through a Google Drive link: https://drive.google.com/file/d/1qUr4buFPajm3bRUvZabjPUtfzIhcyEvM/view?usp=sharing. These resources will be deposited in a public research repository before publication.

## Conflict of Interest Statement

The authors declare no potential conflicts of interest.

## Ethics statement

All participants provided written informed consent for the use of their samples. The use of human-derived PDAC organoid samples was approved by the Ethics Committee of Helsinki University Central Hospital under approval numbers 2075/2016, 658/2019, 2761/2021, and 95/2021. All procedures were conducted in accordance with the Declaration of Helsinki.

## Notes

### Competing Interest Statement

The authors have declared no competing interest.

### Summary of Updates

The revised version addresses reviewer comments through systematic restructuring of the manuscript, including reorganization of the Results and Discussion, revised figures/tables, and clarified presentation of methods and findings. We have also added new analyses such as ablation analysis of scDEAL, bioinformatics analyses towards lineage-tracing datasets, etc. These additions do not alter the main conclusions but strengthen the robustness of the reported findings.

## References

1. Lawson, D.A., et al., Tumour heterogeneity and metastasis at single-cell resolution. Nat Cell Biol, 2018. 20(12): p. 1349–1360.

2. Van de Sande, B., et al., Applications of single-cell RNA sequencing in drug discovery and development. Nature Reviews Drug Discovery, 2023. 22(6): p. 496–520.

3. Maeser, D., et al., A review of computational methods for predicting cancer drug response at the single-cell level through integration with bulk RNAseq data. Curr Opin Struct Biol, 2024. 84: p. 102745.

4. Wu, Z., et al., Single-Cell Techniques and Deep Learning in Predicting Drug Response. Trends Pharmacol Sci, 2020. 41(12): p. 1050–1065.

5. Hao, M., et al., Large-scale foundation model on single-cell transcriptomics. Nat Methods, 2024. 21(8): p. 1481–1491.

6. Cui, H., et al., scGPT: toward building a foundation model for single-cell multi-omics using generative AI. Nat Methods, 2024. 21(8): p. 1470–1480.

7. Wang, Q., et al., scDrugMap: Benchmarking Large Foundation Models for Drug Response Prediction. arXiv e-prints, 2025: p. arXiv:2505.05612.

8. DenAdel, A., et al., Evaluating the role of pre-training dataset size and diversity on single-cell foundation model performance. bioRxiv, 2024: p. 2024.12.13.628448.

9. Yuan, X., et al., Cell ontology guided transcriptome foundation model, in Advances in Neural Information Processing Systems, A. Globerson, et al., Editors. 2024, Curran Associates, Inc. p. 6323–6366.

10. Loftus, T.J., et al., Ideal algorithms in healthcare: Explainable, dynamic, precise, autonomous, fair, and reproducible. PLOS Digit Health, 2022. 1(1).

11. Iorio, F., et al., A Landscape of Pharmacogenomic Interactions in Cancer. Cell, 2016. 166(3): p. 740–754.

12. Barretina, J., et al., The Cancer Cell Line Encyclopedia enables predictive modelling of anticancer drug sensitivity. Nature, 2012. 483(7391): p. 603–607.

13. Xia, F., et al., A cross-study analysis of drug response prediction in cancer cell lines. Briefings in Bioinformatics, 2022. 23(1): p. bbab356.

14. Ovchinnikova, K., et al., Overcoming limitations in current measures of drug response may enable AI-driven precision oncology. npj Precision Oncology, 2024. 8(1): p. 95.

15. Lenhof, K., et al., Simultaneous regression and classification for drug sensitivity prediction using an advanced random forest method. Scientific Reports, 2022. 12(1): p. 13458.

16. Narykov, O., et al., Data imbalance in drug response prediction: multi-objective optimization approach in deep learning setting. Briefings in Bioinformatics, 2025. 26(2): p. bbaf134.

17. Bernett, J., et al., From Hype to Health Check: Critical Evaluation of Drug Response Prediction Models with DrEval. bioRxiv, 2025: p. 2025.05.26.655288.

18. Bohl, M., et al., Domain-adaptation deep learning models do not outperform simple baseline models in single-cell anti-cancer drug sensitivity prediction. bioRxiv, 2026: p. 2026.02.24.707713.

19. Yan, F., Z. Du, and Y.A. Huang, DAGFormer: A graph-based domain adaptation approach for single-cell cancer drug response prediction. PLoS Comput Biol, 2025. 21(12): p. e1013832.

20. Dai, J., et al., Tracing back primed resistance in cancer via sister cells. Nat Commun, 2024. 15(1): p. 1158.

21. Oren, Y., et al., Cycling cancer persister cells arise from lineages with distinct programs. Nature, 2021. 596(7873): p. 576–582.

22. Liu, X., et al., DRMref: comprehensive reference map of drug resistance mechanisms in human cancer. Nucleic Acids Res, 2024. 52(D1): p. D1253–D1264.

23. Fustero-Torre, C., et al., Beyondcell: targeting cancer therapeutic heterogeneity in single-cell RNA-seq data. Genome Med, 2021. 13(1): p. 187.

24. Suphavilai, C., et al., Predicting heterogeneity in clone-specific therapeutic vulnerabilities using single-cell transcriptomic signatures. Genome Med, 2021. 13(1): p. 189.

25. Pellecchia, S., et al., Predicting drug response from single-cell expression profiles of tumours. BMC Med, 2023. 21(1): p. 476.

26. Liu, X., et al., DrugFormer: Graph-Enhanced Language Model to Predict Drug Sensitivity. Adv Sci (Weinh), 2024. 11(40): p. e2405861.

27. Chawla, S., et al., Gene expression based inference of cancer drug sensitivity. Nat Commun, 2022. 13(1): p. 5680.

28. Zheng, Z., et al., Enabling Single-Cell Drug Response Annotations from Bulk RNA-Seq Using SCAD. Adv Sci (Weinh), 2023. 10(11): p. e2204113.

29. Chen, J., et al., Deep transfer learning of cancer drug responses by integrating bulk and single-cell RNA-seq data. Nat Commun, 2022. 13(1): p. 6494.

30. Lei, W., et al., scDR: Predicting Drug Response at Single-Cell Resolution. Genes (Basel), 2023. 14(2).

31. Zhang, W., et al., Integration of Pan-Cancer Cell Line and Single-Cell Transcriptomic Profiles Enables Inference of Therapeutic Vulnerabilities in Heterogeneous Tumors. Cancer Res, 2024. 84(12): p. 2021–2033.

32. Makinen, L., et al., Pancreatic Cancer Organoids in the Field of Precision Medicine: A Review of Literature and Experience on Drug Sensitivity Testing with Multiple Readouts and Synergy Scoring. Cancers (Basel), 2022. 14(3).

33. Boj, S.F., et al., Organoid models of human and mouse ductal pancreatic cancer. Cell, 2015. 160(1-2): p. 324–38.

34. Hao, Y., et al., Integrated analysis of multimodal single-cell data. Cell, 2021. 184(13): p. 3573–3587 e29.

35. Korsunsky, I., et al., Fast, sensitive and accurate integration of single-cell data with Harmony. Nat Methods, 2019. 16(12): p. 1289–1296.

36. Hanzelmann, S., R. Castelo, and J. Guinney, GSVA: gene set variation analysis for microarray and RNA-seq data. BMC Bioinformatics, 2013. 14: p. 7.

37. Aibar, S., et al., SCENIC: single-cell regulatory network inference and clustering. Nat Methods, 2017. 14(11): p. 1083–1086.

38. Qiu, X., et al., Single-cell mRNA quantification and differential analysis with Census. Nat Methods, 2017. 14(3): p. 309–315.

39. Zhang, J., et al., ClusterGVis: An Advanced Visualization and Clustering Tool for Gene Expression Analysis. Genomics Proteomics Bioinformatics, 2026.

40. Domcke, S., et al., Evaluating cell lines as tumour models by comparison of genomic profiles. Nat Commun, 2013. 4: p. 2126.

41. Baek, S., K. Song, and I. Lee, Single-cell foundation models: bringing artificial intelligence into cell biology. Exp Mol Med, 2025. 57(10): p. 2169–2181.

42. Anand, U., et al., Cancer chemotherapy and beyond: Current status, drug candidates, associated risks and progress in targeted therapeutics. Genes Dis, 2023. 10(4): p. 1367–1401.

43. Lenhof, K., et al., Reliable anti-cancer drug sensitivity prediction and prioritization. Sci Rep, 2024. 14(1): p. 12303.

44. Boyd, K., et al. Unachievable region in precision-recall space and its effect on empirical evaluation. in Proceedings of the… International Conference on Machine Learning. International Conference on Machine Learning. 2012.

45. Vasan, N., J. Baselga, and D.M. Hyman, A view on drug resistance in cancer. Nature, 2019. 575(7782): p. 299–309.

46. Donati, G. and B. Amati, MYC and therapy resistance in cancer: risks and opportunities. Mol Oncol, 2022. 16(21): p. 3828–3854.

47. Dias, M.P., et al., Understanding and overcoming resistance to PARP inhibitors in cancer therapy. Nat Rev Clin Oncol, 2021. 18(12): p. 773–791.

48. Passaro, A., et al., Overcoming therapy resistance in EGFR-mutant lung cancer. Nat Cancer, 2021. 2(4): p. 377–391.

49. Gkountakos, A., et al., Identification of Targetable Liabilities in the Dynamic Metabolic Profile of EGFR-Mutant Lung Adenocarcinoma: Thinking beyond Genomics for Overcoming EGFR TKI Resistance. Biomedicines, 2022. 10(2).

50. Aversa, I., et al., Chemoresistance in H-Ferritin Silenced Cells: The Role of NF-kappaB. Int J Mol Sci, 2018. 19(10).

51. Weaver, B.A., How Taxol/paclitaxel kills cancer cells. Mol Biol Cell, 2014. 25(18): p. 2677–81.

52. Gascoigne, K.E. and S.S. Taylor, Cancer cells display profound intra- and interline variation following prolonged exposure to antimitotic drugs. Cancer Cell, 2008. 14(2): p. 111–22.

53. Kang, S.H., et al., Inhibition of MEK with trametinib enhances the efficacy of anti-PD-L1 inhibitor by regulating anti-tumor immunity in head and neck squamous cell carcinoma. Oncoimmunology, 2019. 8(1): p. e1515057.

54. Caunt, C.J., et al., MEK1 and MEK2 inhibitors and cancer therapy: the long and winding road. Nat Rev Cancer, 2015. 15(10): p. 577–92.

55. Zhu, Y., et al., Ensemble transfer learning for the prediction of anti-cancer drug response. Sci Rep, 2020. 10(1): p. 18040.

56. Sinha, S., et al., PERCEPTION predicts patient response and resistance to treatment using single-cell transcriptomics of their tumors. Nat Cancer, 2024. 5(6): p. 938–952.

57. Tasci, E., et al., Bias and Class Imbalance in Oncologic Data-Towards Inclusive and Transferrable AI in Large Scale Oncology Data Sets. Cancers (Basel), 2022. 14(12).

58. Werner de Vargas, V., et al., Imbalanced data preprocessing techniques for machine learning: a systematic mapping study. Knowledge and Information Systems, 2023. 65(1): p. 31–57.

59. Grassberger, C., et al., Patient-Specific Tumor Growth Trajectories Determine Persistent and Resistant Cancer Cell Populations during Treatment with Targeted Therapies. Cancer Res, 2019. 79(14): p. 3776–3788.

60. Heidari, M., et al., Evaluating Single-Cell Perturbation Response Models Is Far from Straightforward. bioRxiv, 2026: p. 2026.02.14.705879.

61. Wang, Z., et al., Drug-tolerant persister cells in cancer: bridging the gaps between bench and bedside. Nat Commun, 2025. 16(1): p. 10048.

62. Sharma, S.V., et al., A chromatin-mediated reversible drug-tolerant state in cancer cell subpopulations. Cell, 2010. 141(1): p. 69–80.

63. Marine, J.C., S.J. Dawson, and M.A. Dawson, Non-genetic mechanisms of therapeutic resistance in cancer. Nat Rev Cancer, 2020. 20(12): p. 743–756.

64. Osorio, D., et al., Drug combination prediction for cancer treatment using disease-specific drug response profiles and single-cell transcriptional signatures. 2025, eLife Sciences Publications, Ltd.

