## Supplemental Methods and Figures for "Predicting Pre-treatment Resistance or Post-treatment Effect? A Systematic Benchmarking of Single-Cell Drug Response Models"

**Additional File: Supplementary Methods and Figures**

### Supplementary Methods

The definition, numerical range, and directionality of the drug-response scores generated by each method are summarized in **Supplementary Table S1**. The drug/therapy coverage of each method across the 26 benchmark datasets, together with the balanced sample sizes available for each dataset, is summarized in **Supplementary Table S2**.

### Beyondcell

Beyondcell is an R-based method to identify tumour cell subpopulations with distinct drug responses from scRNA-seq data. It computes enrichment scores across curated drug signatures to identify drug-responsive cellular subpopulations. In our benchmark, preprocessed Seurat objects were used as input, and drug signature collections were retrieved using the *GetCollection* function. Beyondcell provides two types of drug signature collections: (i) drug perturbation signatures (PSc), derived from the CLUE/LINCS database and reflecting transcriptomic differences between perturbed and unperturbed cells, and (ii) drug sensitivity signatures (SSc), derived from pharmacogenomics resources such as CTRP and GDSC, which captures gene expression patterns associated with drug response.

For each benchmark dataset, we computed Beyondcell scores (BCS) using the *bcScore* function. The BCS estimates the enrichment of every drug signature in the selected collection for each cell in the expression matrix. Higher BCS values indicate greater predicted drug sensitivity, whereas lower values indicate greater resistance. Cells were subsequently classified as sensitive or resistant according to the drug-specific switch point (SP) defined by the method. Because a single drug may be associated with multiple curated Beyondcell signatures, we evaluated all available signatures corresponding to the same drug. For each drug, we selected the signature that achieved the highest AUROC (for balanced datasets, Area Under the Receiver Operating Characteristic) ) or normalized AUPRC (for imbalanced datasets, Area Under the Precision-Recall Curve), and used the prediction scores derived from this selected signature as the final Beyondcell output for performance evaluation.

### CaDRReS-sc

CaDRReS-sc predicts drug response by establishing similarity kernel functions between drugs and cells. The model learns latent representations of drugs and cells in a pharmacogenomic space and optimizes an objective function to project predicted $IC_{50}$ values from different drugs onto a comparable scale, thereby enabling direct comparison of a cell’s relative response across multiple drugs.

In our benchmark, we trained CaDRReS-sc using bulk RNA-seq expression profiles of GDSC cell lines together with their corresponding drug response profiles. The drug response matrix was derived from absolute $IC_{50}$ values estimated across nine dosages using Bayesian sigmoid curve fitting, and served as the ground-truth values for model training. The gene expression matrix was aggregated by gene symbol and *log2* mean fold change normalized. We retained only the genes in the essential gene list provided by the original CaDRReS-sc implementation, and used the resulting expression matrix for kernel construction. The maximum tested drug concentration (*log2_max_conc*) was used as drug-specific information to compute loss-based sample weights via a logistic function. Finally, the cell–cell similarity kernel and the $IC_{50}$-based drug response matrix were jointly used to train the CaDRReS-sc model.

Two model specifications were considered:

1) *cadrres-wo-sample-bias*: equivalent to the baseline CaDRReS model with the sample bias term removed, i.e., without correcting for systematic bias at the cell line level.
2) *cadrres-wo-sample-bias-weight*: in addition to removing sample bias, this setting incorporates two weighting schemes: *ciu* (drug-sample weight), a logistic weight based on drug maximum concentration, and *du* (indication-specific weight), a disease/indication-level factor (set to 1 in this study).

Model optimization was performed in the pharmacogenomic latent space with a dimensionality of 10, using a learning rate of 0.01, and a maximum of 100,000 optimization steps.

In the prediction phase, single-cell RNA-seq expression matrices (TPM) were used as input. The test data were processed in the same manner as the training data, including gene symbol aggregation and *log2* mean fold change normalization. The processed data were then combined with the reference GDSC bulk expression matrix to compute kernel features for the test data. The trained model parameters were then loaded, and the *predict_from_model* function was executed independently for each dataset, producing drug response prediction matrices containing $log_2\ (IC_{50})$ values for each cell. Lower predicted values indicate greater drug sensitivity, whereas higher values indicate greater resistance.

**DREEP**

DREEP is an R package that correlates gene expression with drug response data across hundreds of tumor cell lines. Using these signatures, it computes enrichment scores at the single-cell level to infer the response of each cell to a specific drug. The enrichment score ranges from −1 to 1, with lower values indicating greater predicted sensitivity and higher values indicating greater resistance.

During preprocessing, we first applied a Gene Frequency–Inverse Cell Frequency

(GF-ICF) transformation using the *gficf* package to extract the top relevant genes from each cell as the input matrix, and then passed to the *runDREEP* function for prediction, with parameters setting to *n.markers* = 500 and *gsea* = "simple", and drug signature databases specified as CTRP2 and GDSC. Similar to Beyondcell, the enrichment score matrix output by DREEP may contain multiple signatures corresponding to the same drug. In such cases, we selected the signature achieving the best performance based on AUROC or normalized AUPRC to represent the performance of DREEP on that dataset.

**DrugFormer**

DrugFormer is a graph-enhanced language model that integrates gene expression with prior knowledge through a dual-pathway architecture. A Transformer encoder captures gene-level representations, while a graph attention (GAT) network extracts features from a knowledge graph built on haploinsufficiency and copy-number sensitivity–related gene annotations. The two streams are fused via a gated aggregation module, and the resulting embeddings are used for drug response prediction.

In this benchmark, we followed the original DrugFormer workflow, including knowledge graph construction, dataset conversion and drug response prediction. We first built a gene–token dictionary from the rare copy-number variants (rCNV) scores reported by Collins et al [1]. Haploinsufficiency and copy-number sensitivity-related scores were averaged to compute gene-level scores, which defining edge weights between gene $i$ and gene $j$ as $w_{ij} = 1 - |g_i - g_j|$, where $g_i$ and $g_j$ are the corresponding gene scores. Node features were obtained from the rCNV eigenfeature matrix. The resulting graph data was then used as input to the GAT encoder in DrugFormer.

For dataset conversion, raw count matrices were processed into tokenized inputs consistent with the prebuilt gene–token dictionary. Gene names in each dataset were first mapped to their corresponding tokens while unmapped genes were discarded. For each cell, expressed genes were ranked by expression level, and the ordered list of gene tokens was truncated or padded to a fixed length of 2,048. Each sample was assigned a binary label based on the drug response annotation (sensitive = 1, resistant = 0).

The prediction module was trained and evaluated using a five-fold cross-validation strategy. Tokenized gene expression sequences together with the prebuilt knowledge graph were used as model inputs. Training was performed with AdamW optimizer and cross-entropy loss, and performance was evaluated with multiple metrics including accuracy, precision, recall, and F1 score, with AUROC reported for balanced datasets and normalized AUPRC reported for imbalanced datasets.

**Precily**

Precily integrates representations of cellular transcriptional states with vectorized chemical features of drugs to model their interactions. It maps gene expression profiles into pathway-level embeddings and encodes compounds through learned molecular descriptors. These two modalities are combined within a deep neural network that learns a continuous function relating cell states to drug response. The model output is predicted ln(IC50) values, with lower values indicating greater drug sensitivity and higher values indicating greater resistance.

Since the authors provided pretrained models trained on the CCLE dataset in their GitHub repository, we directly loaded these released models for prediction in our benchmarking workflow. Specifically, we first converted the single-cell count matrix into gene-length–normalized TPM values and applied log-transformation. We then computed pathway activity scores using GSVA based on MSigDB canonical pathways to construct an input feature matrix consistent with the original Precily model design. These pathway features were provided as inputs into the five pretrained models, and the final predicted drug response values were obtained by averaging the outputs across all models, as defined in the original implementation.

**SCAD**

SCAD aims to transfer drug response knowledge from bulk cell lines to single-cell data via adversarial domain adaptation. The model architecture integrates a shared feature extractor, a domain discriminator, and a drug response predictor to align latent feature distributions between domains while minimizing prediction error. The drug-response predictor outputs a continuous sensitivity probability between 0 and 1, with higher values indicating greater predicted sensitivity.

We benchmarked SCAD using a unified workflow applied consistently across all the datasets, drugs, and experimental scenarios. For each single-cell dataset, raw count matrices were exported from *Seurat* and processed using the *Scanpy* pipeline, including normalization, log-transformation, and z-score scaling. Following the original SCAD design, we constructed three alternative input feature sets before model training: "*all*", including all genes shared between the source and target domains; "*tp4k*", the top 4,000 highly variable genes identified by *Scanpy*; and "*PPI*", a curated subset of 2,128 protein–protein interaction genes previously reported to be informative for drug response prediction.

Processed single-cell features were aligned with bulk GDSC expression and drug-response profiles to construct matched source-domain (GDSC) and target-domain (single-cell) matrices for each drug. For each dataset–drug pair, SCAD's default stratified five-split scheme was applied: GDSC samples were partitioned into source-training and source-validation sets, while single-cell profiles were divided into target-training and held-out target-test sets. Both imbalance-handling strategies implemented in SCAD were evaluated independently, including (i) weighted sampling and (ii) SMOTE-based over-/undersampling of source-domain training samples.

Using the hyperparameter configuration that achieved the highest average validation AUROC during random search, SCAD was retrained and tested separately on each of the five splits. The random search covered the hidden-layer dimension, latent dimension, number of training epochs, domain-adaptation weight, and source/target mini-batch sizes. Candidate configurations were evaluated across the five source-domain validation splits, and the mean source-domain validation AUROC was used to select the final configuration before target-domain prediction. Performance metrics were computed for each split from the predicted values, and the mean across splits was reported as the final performance.

**scDEAL**

scDEAL predicts single-cell drug responses through a deep transfer learning framework that bridges bulk and single-cell transcriptomic data. Two denoising autoencoders independently extract low-dimensional representations from bulk and single-cell gene expression profiles, and a predictor is trained to model bulk gene–drug response relations. The framework then jointly optimizes these modules to minimize distribution difference between bulk and single-cell features while maintaining accurate bulk-level prediction. The final model outputs a drug-sensitivity probability between 0 and 1 for each cell, with higher values indicating greater predicted sensitivity.

We followed the official implementation of the method for all evaluations. Specifically, we used the integrated and labeled GDSC and CCLE datasets as the bulk-level transcriptomic data for model training. To maintain model stability, we fixed the core architectural parameters according to the authors' recommendations. These settings included using denoising autoencoder for dimensionality reduction, a regularization mode that integrates cell-type information, and fixed hidden dimensions of the Encoder and Predictor set to [512, 256] and [256, 128], respectively. The learning rate and dropout rate were set to 0.5 and 0.1. Building on this configuration, we implemented a systematic hyperparameter grid search covering a wide range of bottleneck dimensions from 8 to 512 (8, 16, 32, 64, 128, 256, 512), and tested three sampling techniques, including up sampling, down sampling, and SMOTE, for handling source domain data. The final single-cell drug response predictions were generated using the transferred models obtained under these settings.

**scDr**

scDr predicts cell-level drug responses by leveraging pharmacogenomic data to derive drug-specific gene signatures. For each drug, resistant and sensitive cell lines are first identified based on their AUROC values in the CTRP dataset. Differential expression analysis between these groups is then performed to identify the top genes as drug-response genes (DRGs). Using single-cell transcriptomic data, scDr calculates a drug-response score (DRS) for each cell.

We first processed the CCLE bulk RNA-seq expression profiles and CTRP drug response data to establish drug-specific signatures. For each compound, resistant and sensitive cell lines were identified based on their AUROC distributions, followed by differential expression analysis to determine the drug-response signatures. These signatures were then applied to single-cell datasets to compute drug-response scores. For each drug, the normalized expression of its signature genes in the scRNA-seq dataset was weighted by the direction of fold change derived from bulk analysis, and the resulting aggregated score represented the drug-response score of each cell, with higher values indicating greater predicted resistance and lower values indicating greater sensitivity.

**scIDUC**

Similar to scDr, scIDUC predicts cell-level drug responses by identifying drug-response genes (DRGs) for each compound and integrating bulk and single-cell transcriptomic data into a shared latent space using algorithms such as canonical correlation analysis (CCA) or non-negative matrix factorization (NMF). Within this shared representation, regression-based models are applied to infer drug response at the single-cell level. The resulting continuous predictions represent estimated normalized drug-response AUC values, with lower values indicating greater sensitivity and higher values indicating greater resistance.

As the source code for scIDUC is not publicly available, all prediction tasks were performed using its RShiny-based web application (https://oncotherapyinformatics.org/sciduc_shiny/). To ensure reproducibility and efficiency, an automated workflow was implemented using the Selenium library. This pipeline programmatically handled data uploading, selection of the reference bulk RNA-seq dataset, drug selection, model execution, and automatic retrieval of prediction results.

**PDAC organoid culture and drug treatment**

PDAC organoid lines (PO83T and PO84T) were established and characterized as previously described [2]. Organoids were cultured in Matrigel (Corning) domes and maintained in complete feeding medium following the protocol by Boj et al [3]. For the transcriptional profiling of drug responses, 20,000 dissociated tumor organoid cells from PO83T and PO84T were seeded in 200 µl of a feeding medium/Matrigel (4 mg/ml final) mixture in 24-well plates. The Matrigel mixture was allowed to solidify for 30 minutes at 37°C, after which each well was topped up with 500 µl of feeding medium. On day 4, 50% of the medium was replaced with fresh feeding medium, and the organoids were exposed to 12.5 nM trametinib (MedChemExpress EU, #HY-10999), 500 nM paclitaxel (MedChemExpress EU, # HY-B0015), or a combination of both for 72 hours. DMSO (Merck, #D4540) was used as a vehicle control.

**Single-cell dissociation and cell hashing**

After 72 hours of drug exposure, the culture medium was removed, and the Matrigel domes were gently washed twice with PBS. To generate a single-cell suspension, the domes and organoids were sequentially digested using 250 µl Dispase II (#SCM133, Sigma-Aldrich, #SCM133) and 250 µl TrypLE Express (Gibco/Thermo Fisher Scientific, #12605010), each at 37°C for 20 minutes. Digestion was neutralized by adding a feeding medium supplemented with 10% FBS, and the cells were washed three times with 1x PBS (300 × g, 5 minutes). Cell viability and concentration were assessed using the LUNA-FX7™ Automated Cell Counter (Logos Biosystems, Inc.).

For sample multiplexing, single cells were labeled with TotalSeq™-A anti-human Hashtag antibodies (BioLegend; Cat. Nos. 394601, 394603, and 394605; RRIDs: AB_2750015, AB_2750016, and AB_2750017, respectively) according to the manufacturer's instructions. Labeled cells were resuspended in 1x PBS containing 0.04% BSA at a final concentration of 700–1,200 cells/µl.

**scRNA-seq library preparation and sequencing**

Subsequently, approximately 2,500 to 10,000 cells per sample were loaded into the 10x Genomics Chromium Next GEM Automated system. Single-cell 3' gene expression libraries were constructed using the Chromium Next GEM Automated SC 3' Reagent Suite v3.1 (10x Genomics, #PN-1000121). The final libraries were pooled and sequenced on a NovaSeq SP (Illumina) platform.

**Single-cell RNA-seq data preprocessing and integration**

The scRNA-seq dataset of PDAC organoids was loaded and processed using the R package Seurat (version 4.3.0, RRID:SCR_016341) [4]. Data normalization was performed and highly variable features were identified for each sample individually using the variance-stabilizing transformation method, selecting the top 2,000 features. To account for batch effects across different samples, integration features were determined using SelectIntegrationFeatures. Following data scaling, Principal Component Analysis (PCA) was conducted on the integration features. We subsequently applied the Harmony algorithm via the harmony R package to correct batch effects [5], utilizing the first 30 principal components with the parameters theta = 6 and lambda = 0.3. Unsupervised cell clustering was performed using the FindNeighbors and FindClusters functions with a resolution of 0.5.

**Identification of drug resistance signatures and functional enrichment**

Based on the predicted drug sensitivity probabilities, cells were stratified into "sensitive" and "resistant" populations using the median prediction probability as the threshold. To eliminate confounding effects from cell cycle heterogeneity, cell cycle phases were scored using canonical S and G2/M phase markers, and these scores were subsequently regressed out during data scaling.

Differential gene expression analysis between predicted resistant and sensitive cells was conducted via Seurat's FindMarkers function. To quantify MoA-relevant pathway activities at single-cell resolution, single-sample GSEA (ssGSEA) was implemented using the GSVA package [6]. Besides, custom gene signatures, including drug resistance-specific pathways and MAPK feedback targets, were scored for individual cells utilizing the AUCell algorithm [7].

**Single-Cell trajectory and dynamic gene expression analysis**

Pseudotime trajectory analysis was performed using the monocle R package (version 2.34.0, RRID:SCR_018685) to reconstruct the evolutionary dynamics of drug resistance [8]. The raw count matrix was used to create a CellDataSet object and size factors and dispersions were estimated according to the standard Monocle 2 pipeline. To order cells along the pseudotime trajectory, we selected the top 300 and 1,000 significant differentially expressed genes (DEGs) for the paclitaxel and trametinib datasets, respectively, ranked by their absolute log2-fold changes between the sensitive and resistant populations. Dimensionality reduction was performed using the Discriminative Dimensionality Reduction with Trees (DDRTree) algorithm with a maximum of two components. To explore genes whose expression dynamically changed across cell states along the trajectory, we performed differential expression analysis across all expressed genes to identify state-dependent genes. Genes with a q-value < 1*10-10 were defined as significantly associated with the resistance transition. Finally, these significant dynamic genes were clustered and functionally enriched along the pseudotime trajectory utilizing the ClusterGVis package [9].

**Foundation model benchmarking**

To compare conventional drug-response prediction methods with recent foundation models, we evaluated scGPT and scFoundation using the same balanced, imbalanced and lineage-tracing datasets generated for the original benchmark. For both foundation models, the pretrained models were used to generate cell-level embeddings, which were then used as input to a multilayer perceptron classifier for drug-response prediction. In the within-dataset setting, each dataset was evaluated using stratified three-fold cross-validation, and out-of-fold predictions from the three folds were combined to calculate the final performance. The same within-dataset procedure was applied to the lineage-tracing datasets using their corresponding drug-response labels.

We further evaluated cross-dataset prediction for drugs represented by multiple single-cell datasets. For each drug, the classifier was trained using cell embeddings from one dataset and directly applied to another dataset without using cells from the test dataset during training. All available directional train–test combinations were evaluated. The same pretrained embeddings and downstream classifier settings were used in the within- and cross-dataset analyses.

Performance was evaluated using the same metrics as in the original benchmark. scDEAL results from the corresponding dataset and evaluation settings were used as the conventional reference. For dataset-level comparisons, only datasets shared by scDEAL, scGPT, and scFoundation were retained. When a target dataset had multiple cross-dataset predictions from different training datasets, performance was first averaged across the training datasets within each sampling round and then averaged across the three rounds. Performance differences between each foundation-model setting and scDEAL were calculated as:

$$\Delta_{performance} = Performance_{FM} - Performance_{scDEAL}$$

**Bioinformatics analysis of lineage-tracing datasets**

Transcriptomic differences between pre-sensitive and pre-resistant cells were analyzed in both ReSisTrace and Watermelon. Cell-cycle scores were calculated using canonical S- and G2/M-phase gene sets and included as covariates in differential expression analysis to reduce the influence of cell-cycle variation. Differentially expressed genes were identified using MAST. For visualization and signal-strength comparisons, DEGs were defined using $P < 0.05$ and an absolute log2 fold change > 0.1; FDR-adjusted significance was also calculated. For Watermelon post-treatment samples, replicate effects were corrected using Harmony before UMAP visualization.

Gene set enrichment analysis was performed using Hallmark, KEGG, and Gene Ontology molecular-function gene sets from MsigDB [10, 11]. Genes were ranked by the direction of the fold change weighted by statistical significance, using sign(log2FC) × −log10(P), and enrichment was calculated with clusterProfiler [12]. Pathways with $P < 0.05$ were used for visualization. To compare the strength of pre-existing and treatment-associated transcriptional signals, we summarized DEG numbers and absolute log2 fold changes for each contrast. In ReSisTrace, the intrinsic pre-sensitive versus pre-resistant contrast was compared with the conventional pre-versus post-treatment contrast. In Watermelon, the intrinsic D0 contrast was compared with changes between D0 pre-resistant cells and their resistant counterparts at D3, D7, and D14.

### Supplementary Tables

| Method | Model output | Interpretation | Score range | Binary cutoff | Direction |
|---|---|---|---|---|---|
| **Beyondcell** | Beyondcell Score (BCS) | Enrichment score reflecting the association of each cell with a drug-specific signature. For drug sensitivity signatures, BCS represents predicted drug sensitivity. | Continuous | Drug-specific switch point (SP) | ↑: more sensitive<br>↓: more resistant |
| **CaDRReS-sc** | Predicted log2(IC50) | Predicted half-maximal inhibitory concentration derived from the learned pharmacogenomic latent space. | Continuous | Dataset-specific median of predicted values | ↑: more resistant<br>↓: more sensitive |
| **DREEP** | Enrichment Score (ES) | GSEA-based enrichment of cell-specific relevant genes against the drug-specific genomic profile of drug sensitivity. | −1 to 1 | 0 | ↑: more resistant<br>↓: more sensitive |
| **DrugFormer** | Drug-response probability | Predicted probability for the sensitive or resistant response class generated by the classification model. | 0–1 | Argmax of the resistant and sensitive class outputs | ↑: more sensitive<br>↓: more resistant |
| **Precily** | Predicted ln(IC50) | Predicted drug-response value based on pathway-level cellular features and drug molecular descriptors. | Continuous | Dataset-specific median of predicted values | ↑: more resistant<br>↓: more sensitive |
| **SCAD** | Drug-sensitivity probability | Probability of drug sensitivity predicted after adversarial domain adaptation between bulk and single-cell data. | 0–1 | Dataset-specific median of predicted values | ↑: more sensitive<br>↓: more resistant |
| **scDEAL** | Drug-sensitivity probability | Probability that an individual cell is sensitive to the evaluated drug. | 0–1 | Argmax of the resistance and sensitivity outputs | ↑: more sensitive<br>↓: more resistant |
| **scDr** | Drug Response Score (DRS) | Weighted expression score derived from drug-response genes identified from pharmacogenomic data. | Continuous | Dataset-specific median of predicted values | ↑: more resistant<br>↓: more sensitive |
| **scIDUC** | Predicted normalized AUC (nAUC) | Regression-based estimate of drug response derived from dose-response viability information. | Continuous | Dataset-specific median of predicted values | ↑: more resistant<br>↓: more sensitive |

**Table S1. Definition and interpretation of drug-response scores generated by the evaluated methods.**

| **Dataset** | **Drug Therapy** | **Balanced sample sizes** | **Beyondcell** | **CaDRReS-sc** | **DREEP** | **DrugFormer** | **Precily** | **SCAD** | **scDEAL** | **scDr** | **scIDUC** |
|---|---|---|---|---|---|---|---|---|---|---|---|
| GSE104987 | KDM5-C70 | 500, 1000, 2000 | U | U | U | ✓ | U | U | U | U | U |
| GSE108397 | PLX-4720 | 500, 1000, 2000, 3000, 5000 | ✓ | ✓ | ✓ | ✓ | ✓ | ✓ | ✓ | ✓ | ✓ |
| GSE111014 | Ibrutinib | 500, 1000, 2000, 3000, 5000 | ✓ | U | ✓ | ✓ | ✓ | U | ✓ | ✓ | ✓ |
| GSE115251 | TAE684 | 500, 1000, 2000, 3000, 5000 | ✓ | ✓ | ✓ | ✓ | U | ✓ | U | U | ✓ |
| GSE131984_JQ1 | JQ1 | 500, 1000 | U | ✓ | ✓ | ✓ | ✓ | ✓ | U | U | ✓ |
| GSE131984_Pac | Paclitaxel | 500, 1000 | ✓ | ✓ | ✓ | ✓ | ✓ | ✓ | ✓ | ✓ | ✓ |
| GSE131984_Pal | Palbociclib | 500, 1000 | ✓ | ✓ | ✓ | ✓ | ✓ | ✓ | ✓ | U | ✓ |
| GSE139386 | Ceritinib | 500, 1000, 2000, 3000 | U | U | U | ✓ | U | U | U | U | U |
| GSE140440 | Docetaxel | 1000 | ✓ | ✓ | ✓ | ✓ | ✓ | ✓ | ✓ | ✓ | ✓ |
| GSE149214 | Erlotinib | 500, 1000, 2000 | ✓ | ✓ | ✓ | ✓ | ✓ | ✓ | ✓ | ✓ | ✓ |
| GSE152469 | Ibrutinib | 500, 1000, 2000, 3000, 5000 | ✓ | U | ✓ | ✓ | ✓ | U | ✓ | ✓ | ✓ |
| GSE153697 | anti-CD19 CAR-T | 500, 1000, 2000 | U | U | U | ✓ | U | U | U | U | U |
| GSE158457 | Dasatinib | 500, 1000, 2000, 3000, 5000 | ✓ | ✓ | ✓ | ✓ | ✓ | ✓ | ✓ | ✓ | ✓ |
| GSE162117 | Ficlatuzumab | 500, 1000, 2000, 3000, 5000 | U | U | U | ✓ | U | U | U | U | U |
| GSE163836 | Paclitaxel | 500, 1000, 2000, 3000 | ✓ | ✓ | ✓ | ✓ | ✓ | ✓ | ✓ | ✓ | ✓ |

| | | | | | | | | | | | |
|---|---|---|---|---|---|---|---|---|---|---|---|
| GSE164237 | Pembrolizumab | 500, 1000, 2000, 3000, 5000 | U | U | U | ✓ | U | U | U | U | U |
| GSE164551 | CAR-T | 500, 1000, 2000, 3000, 5000 | U | U | U | ✓ | U | U | U | U | U |
| GSE164897 | Vemurafenib | 500, 1000, 2000, 3000, 5000 | ✓ | U | U | ✓ | U | U | U | U | ✓ |
| GSE168668 | Enzalutamide | 500, 1000, 2000, 3000 | ✓ | U | U | ✓ | U | U | U | U | U |
| GSE169246_Blood | Paclitaxel | 500, 1000, 2000, 3000, 5000 | ✓ | ✓ | ✓ | ✓ | ✓ | ✓ | ✓ | ✓ | ✓ |
| GSE169246_Tissue | Paclitaxel | 500, 1000, 2000, 3000, 5000 | ✓ | ✓ | ✓ | ✓ | ✓ | ✓ | ✓ | ✓ | ✓ |
| GSE175716 | Sorafenib | 500, 1000 | ✓ | ✓ | ✓ | ✓ | ✓ | ✓ | ✓ | ✓ | ✓ |
| GSE186960 | Gemcitabine | 500, 1000 | ✓ | ✓ | ✓ | ✓ | ✓ | ✓ | ✓ | ✓ | ✓ |
| GSE199333 | Gilteritinib | 500, 1000, 2000, 3000, 5000 | U | U | U | ✓ | U | U | U | U | U |
| GSE212217 | Pembrolizumab | 500, 1000, 2000, 3000, 5000 | U | U | U | ✓ | U | U | U | U | U |
| PMID34715028 | Nivolumab | 500, 1000, 2000, 3000, 5000 | U | U | U | ✓ | U | U | U | U | U |

**Table S2. Drug/therapy coverage of the evaluated methods across the benchmark datasets.** Each row represents one benchmark dataset and its corresponding drug or therapy. Check marks (✓) indicate that the drug or therapy is supported by the method, whereas U indicates that it is unsupported. “Balanced sample sizes” indicates the total cell numbers that could be constructed for balanced evaluation based on the available sensitive and resistant cells in each dataset.

### Supplementary Figures

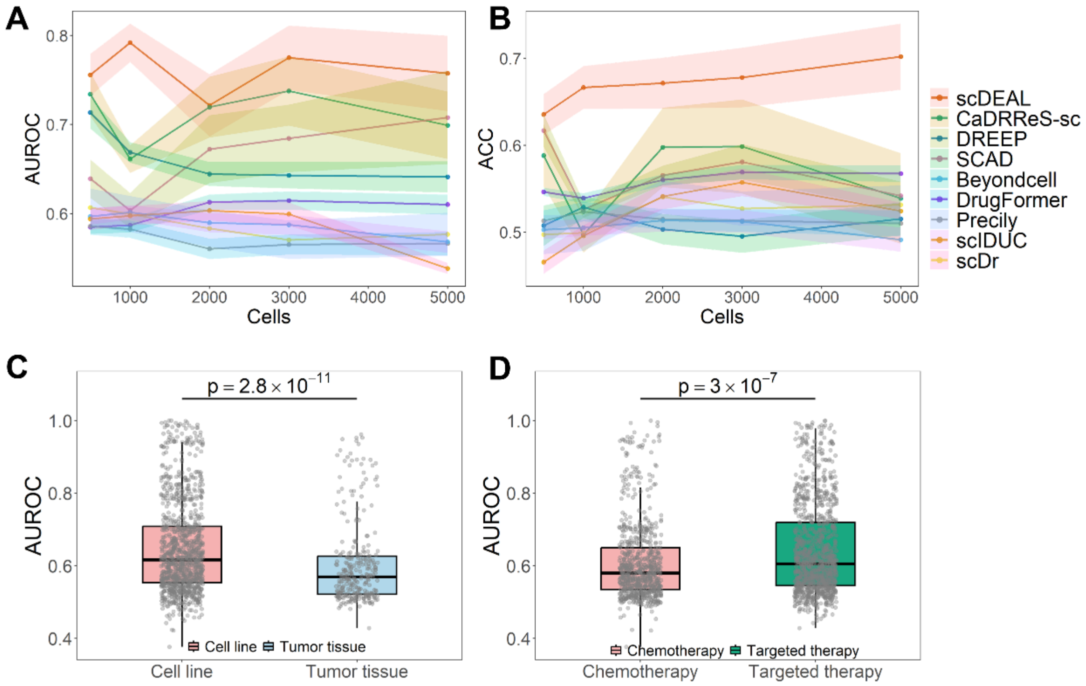


**Figure S1. Additional performance analyses on balanced datasets. (A, B)** Trends of mean AUROC (A) and accuracy (B) with increasing total cell numbers. **(C, D)** Overall AUROC comparisons grouped by sample source (**C**) and therapy type (**D**).

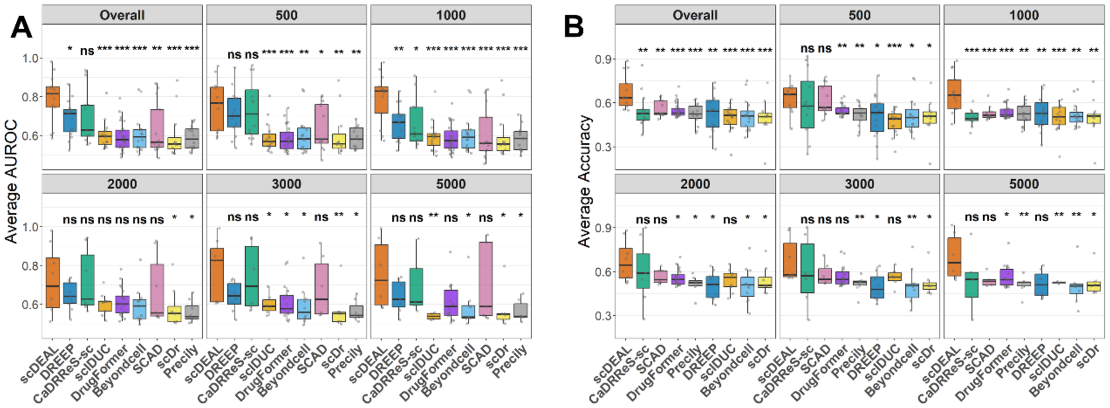

**Figure S2. Performance of scDEAL and other methods across varying training sample sizes (500–5000 cells). (A)** Average AUROC; **(B)** Average accuracy. *P < 0.05, **P < 0.01, ***P < 0.001, and ****P < 0.0001; ns, not significant.

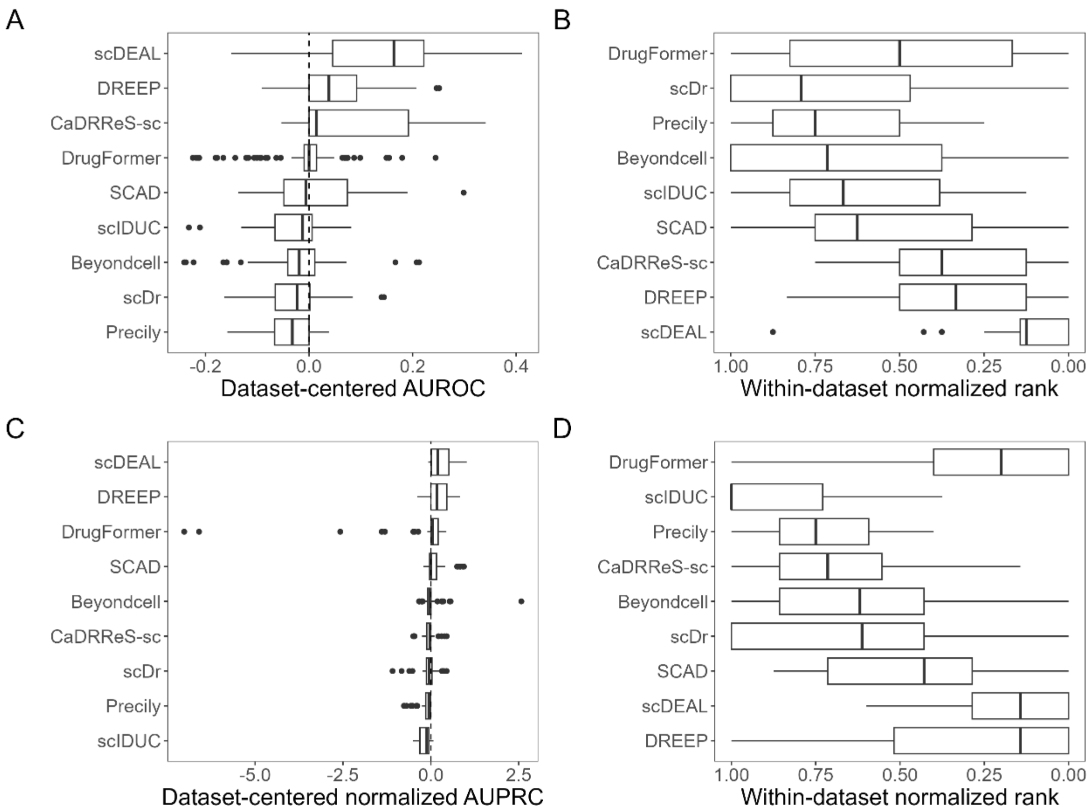


**Figure S3. Within-dataset comparison of model performance.** (**A, B**) Dataset-centered performance (**A**) and within-dataset normalized rank (**B**) of the evaluated methods under balanced settings. (**C, D**) Dataset-centered performance (**C**) and within-dataset normalized rank (**D**) under imbalanced settings. Dataset-centered performance was measured by AUROC for balanced datasets and normalized AUPRC for imbalanced datasets and was calculated by subtracting the median performance across methods within each dataset and sampling setting. Ranks were normalized from 0 to 1, with lower values indicating better relative performance.

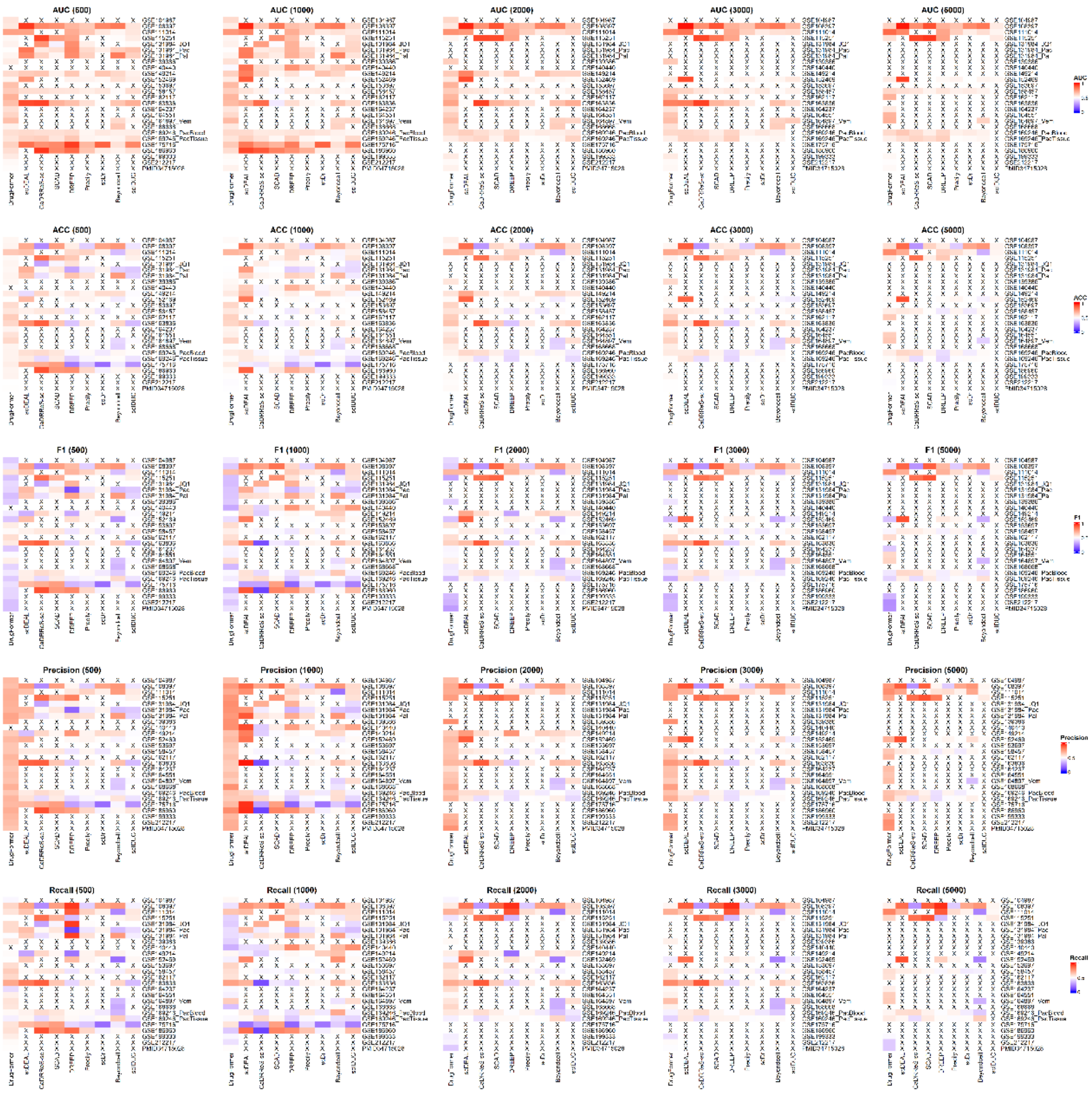

**Figure S4. Dataset-level evaluation across diverse sample sizes.** Heatmaps (top to bottom) showing the AUC, ACC, F1-score, Precision, and Recall for the benchmarked methods across five sample sizes. Each row represents a specific dataset, and columns denote computational tools. Values represent the mean of three independent experimental rounds. Crosses (×) indicate cases where a method failed to generate predictions.

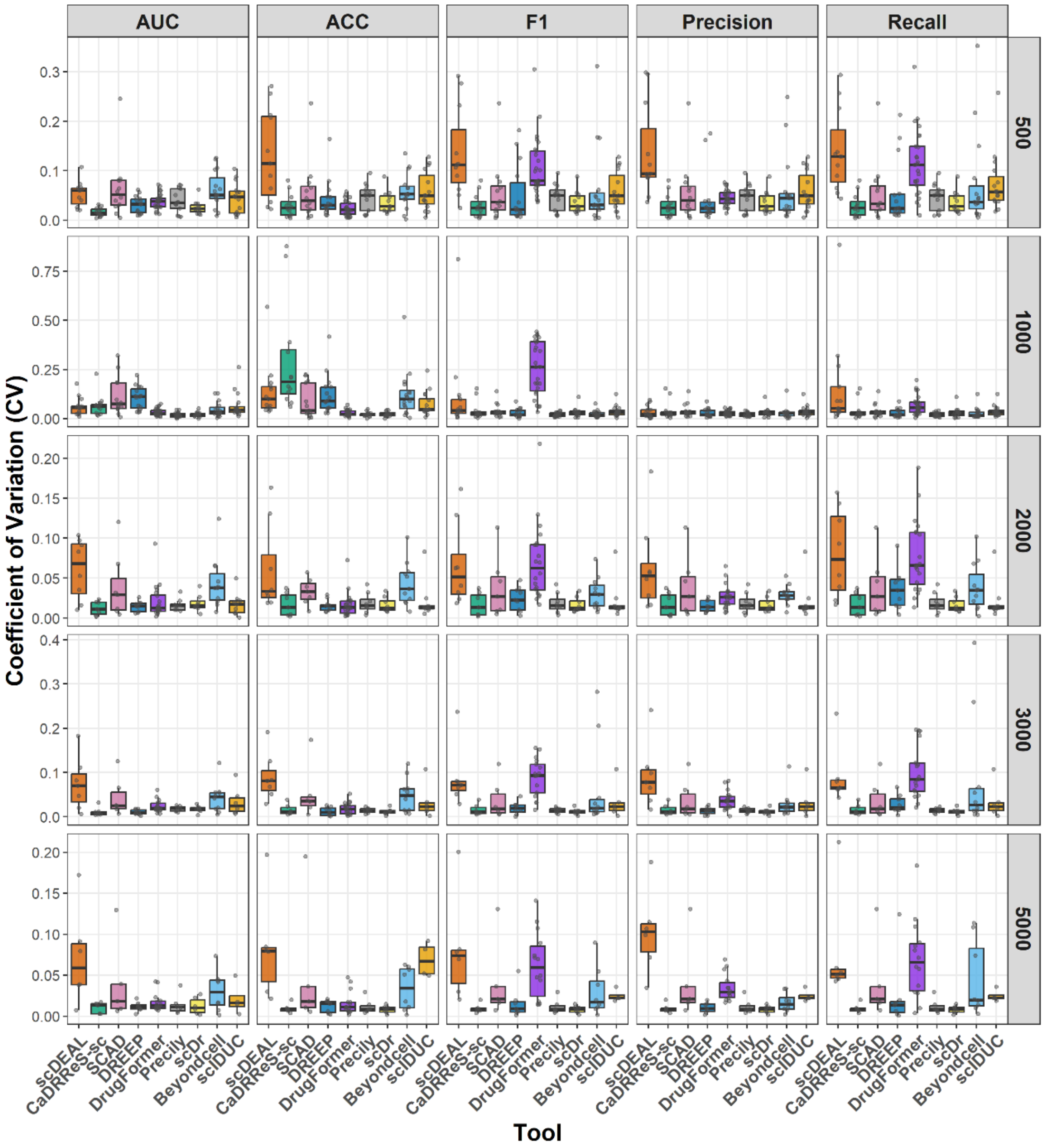


**Figure S5. Stability analysis across different sample sizes.** The coefficient of variation (CV) for five performance metrics is shown at sample sizes ranging from 500 to 5,000 cells. Lower CV indicates higher greater consistency across runs.

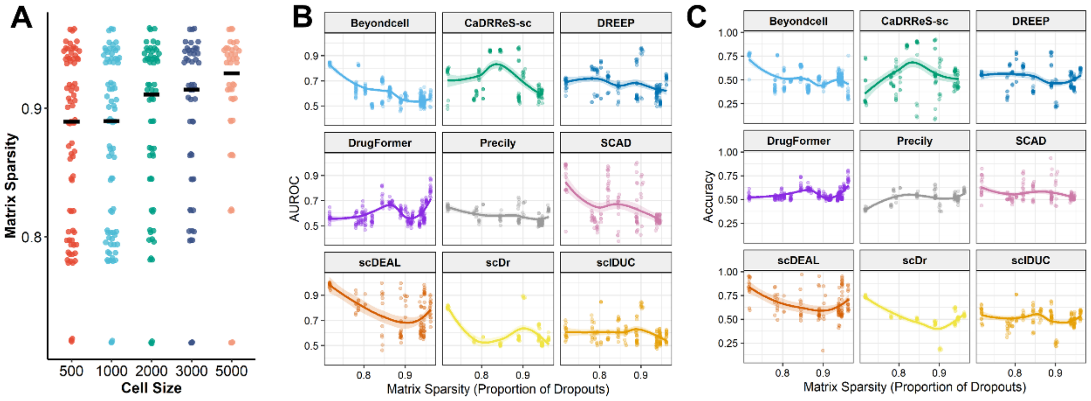


**Figure S6. Association between matrix sparsity and predictive performance. (A)** Distribution of matrix sparsity across subsampled cell sizes ranging from 500 to 5000 cells. **(B–C)** Relationships between matrix sparsity and model performance measured by AUROC **(B)** and Accuracy **(C)** across nine methods.

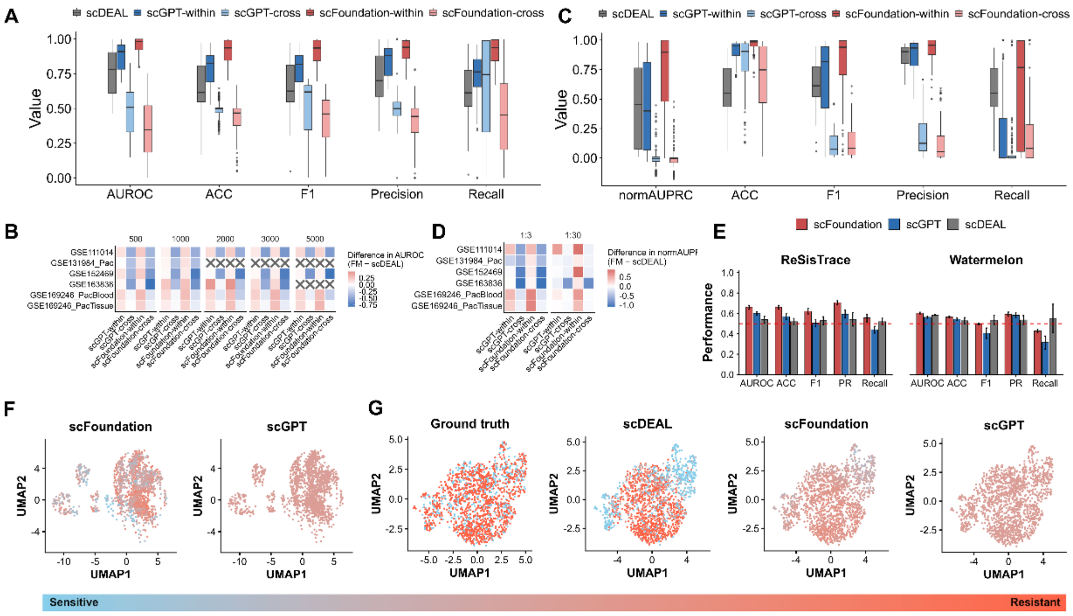


**Figure S7. Evaluation of foundation models across balanced, imbalanced, and lineage-tracing settings. (A)** Performance comparison of scDEAL, scGPT, and scFoundation on balanced datasets, with within-dataset and cross-dataset settings shown separately for scGPT and scFoundation. **(B)** Differences in AUROC between foundation models and scDEAL across datasets and cell numbers in the balanced setting. Crosses indicate unavailable comparisons. (**C**) Performance comparison under imbalanced settings. **(D)** Differences in normalized AUPRC between foundation models and scDEAL under 1:3 and 1:30 class ratios. **(E)** Performance comparison of scFoundation, scGPT, and scDEAL on the ReSisTrace and Watermelon lineage-tracing datasets. **(F)** UMAP visualization of predicted drug-response scores from scFoundation and scGPT in the ReSisTrace dataset. **(G)** UMAP visualization of ground-truth labels and predicted drug-response scores from scDEAL, scFoundation, and scGPT in the Watermelon dataset.

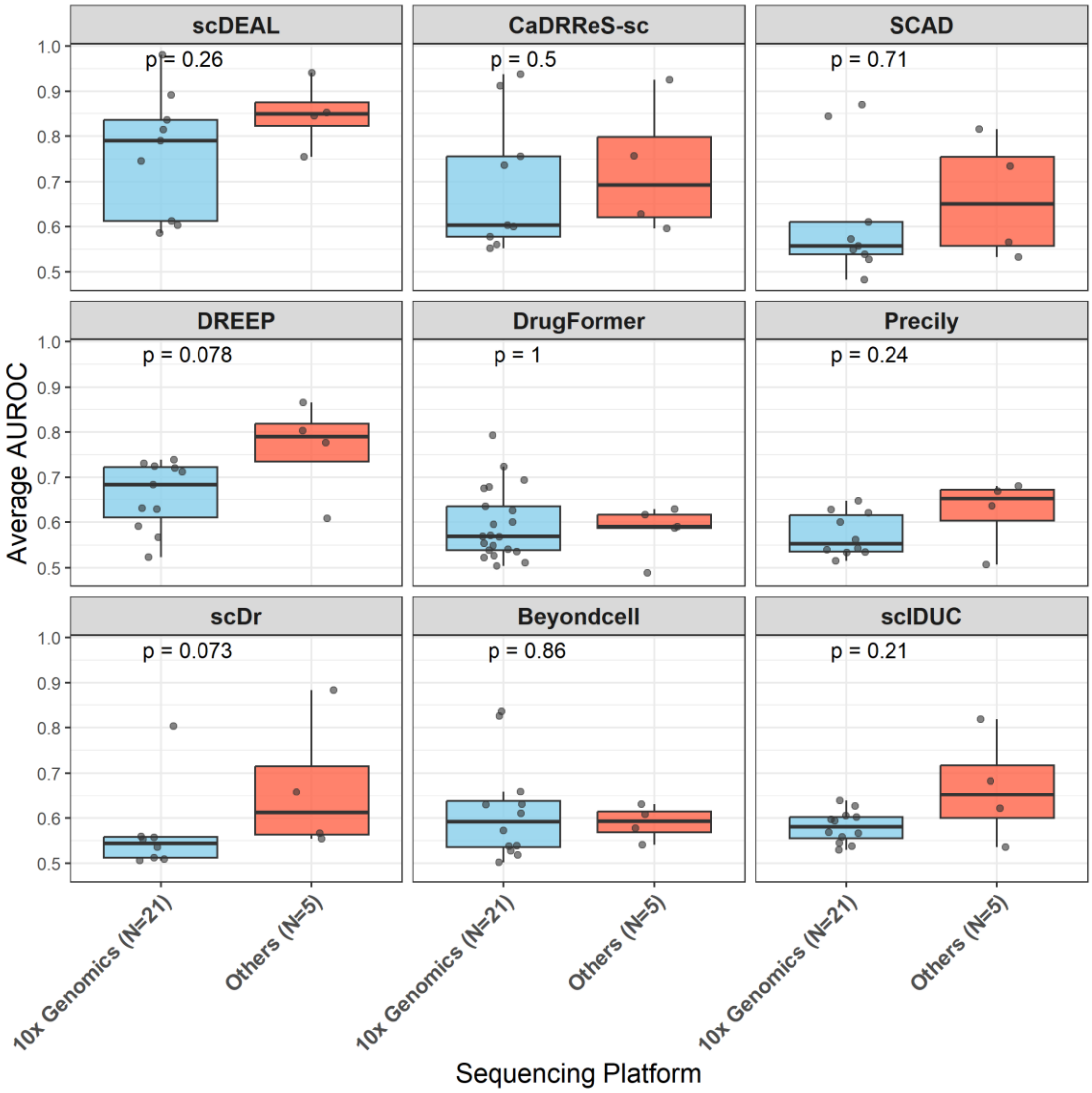


**Figure S8. AUROC comparison between the 10x Genomics platform and other sequencing platforms.**

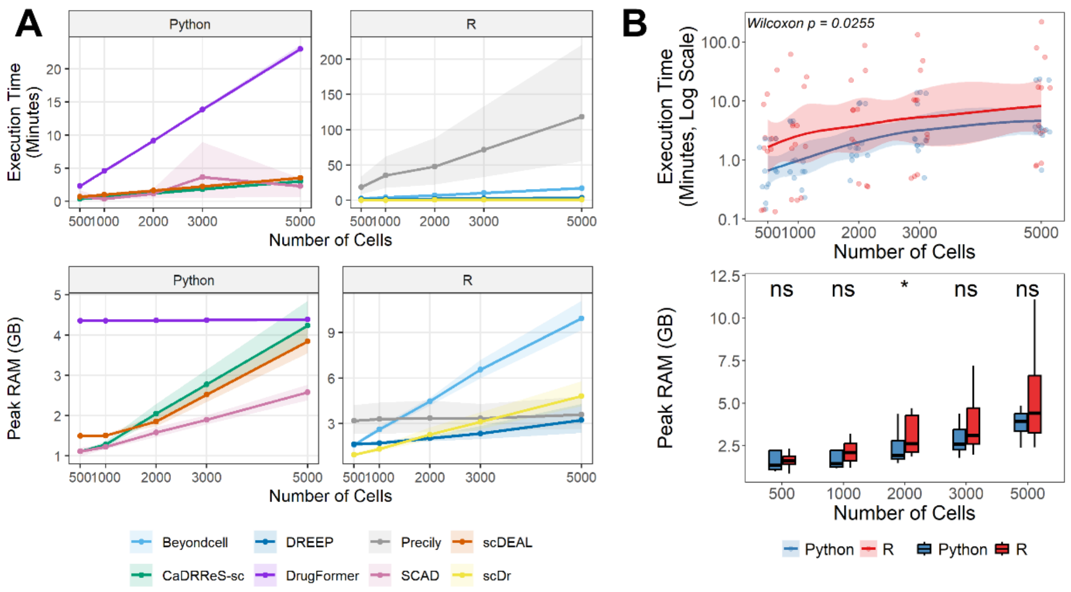


**Figure S9. Benchmarking of computational resource demands across methods. (A)** Execution time and peak RAM usage of different methods across increasing sample size. **(B)** Comparison of Python- and R-based methods in execution time and memory usage.

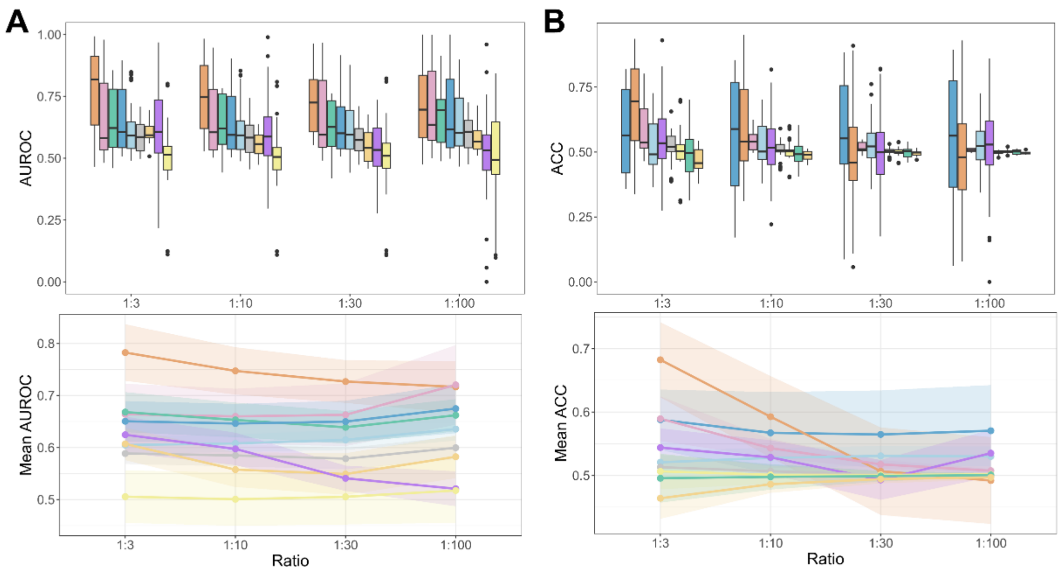


**Figure S10. Additional performance analyses on imbalanced datasets. (A)** AUROC distributions (top) and mean AUROC trends (bottom) across different imbalance ratios. **(B)** ACC distributions (top) and mean ACC trends (bottom) across different imbalance ratios.

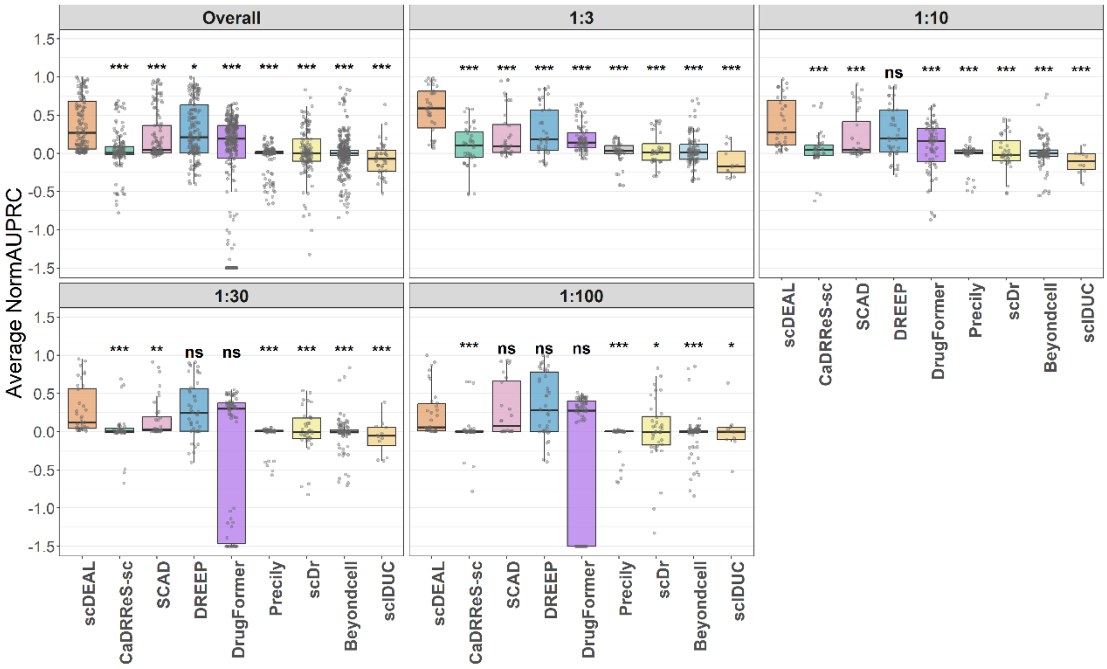


**Figure S11. normAUPRC comparison under class imbalance.** Average normAUPRC of nine methods across different imbalance ratios (1:3, 1:10, 1:30, and 1:100). *P < 0.05, **P < 0.01, ***P < 0.001, and ****P < 0.0001; ns, not significant.

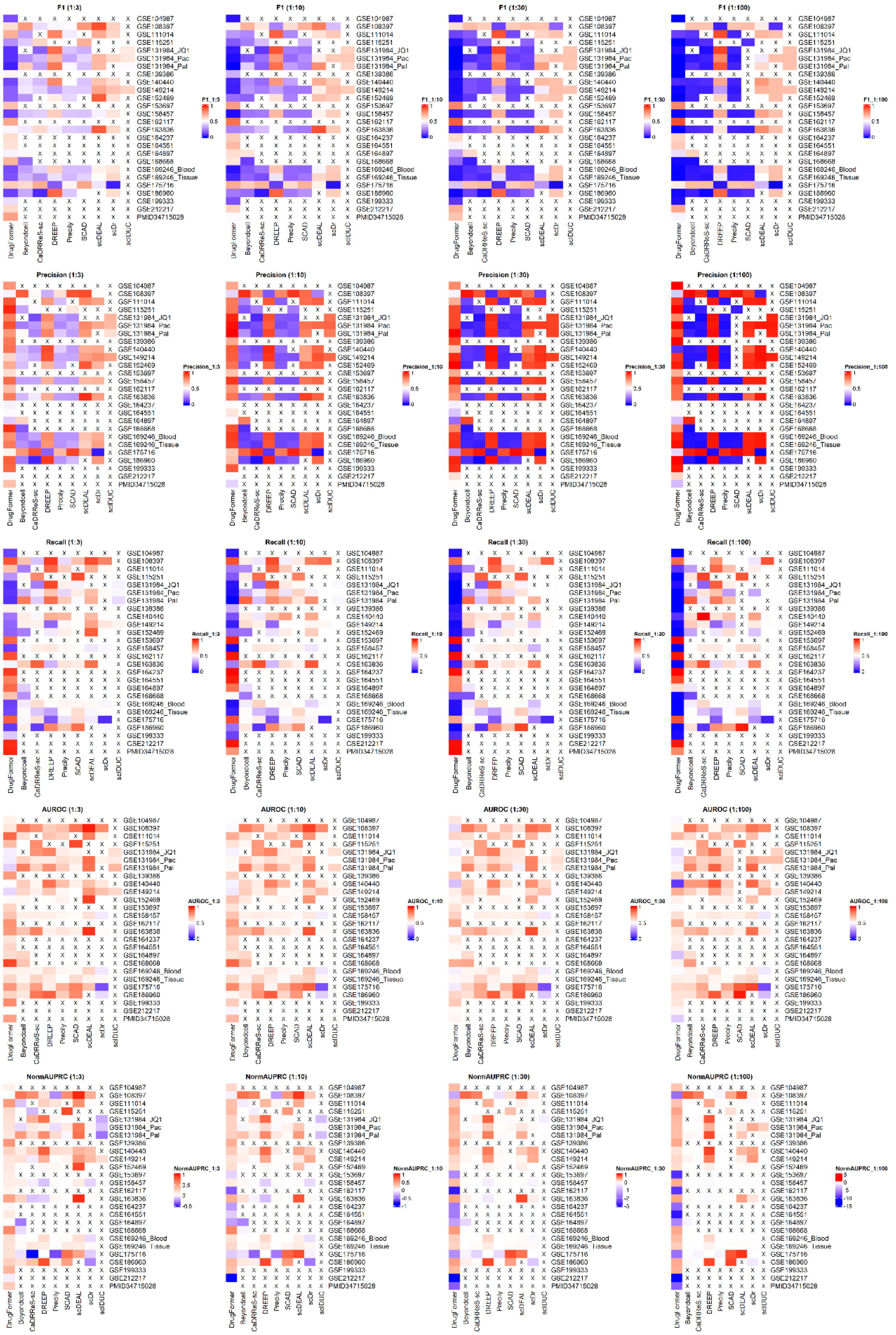


**Figure S12. Dataset-level evaluation across diverse imbalance ratios.** Heatmaps showing the F1-score (top), Precision (middle), and Recall (bottom) for the benchmarked methods across four imbalance ratios. Each row represents a specific dataset, and columns denote computational tools. Values represent the mean of three independent experimental rounds. Crosses (×) indicate cases where a method failed to generate predictions.

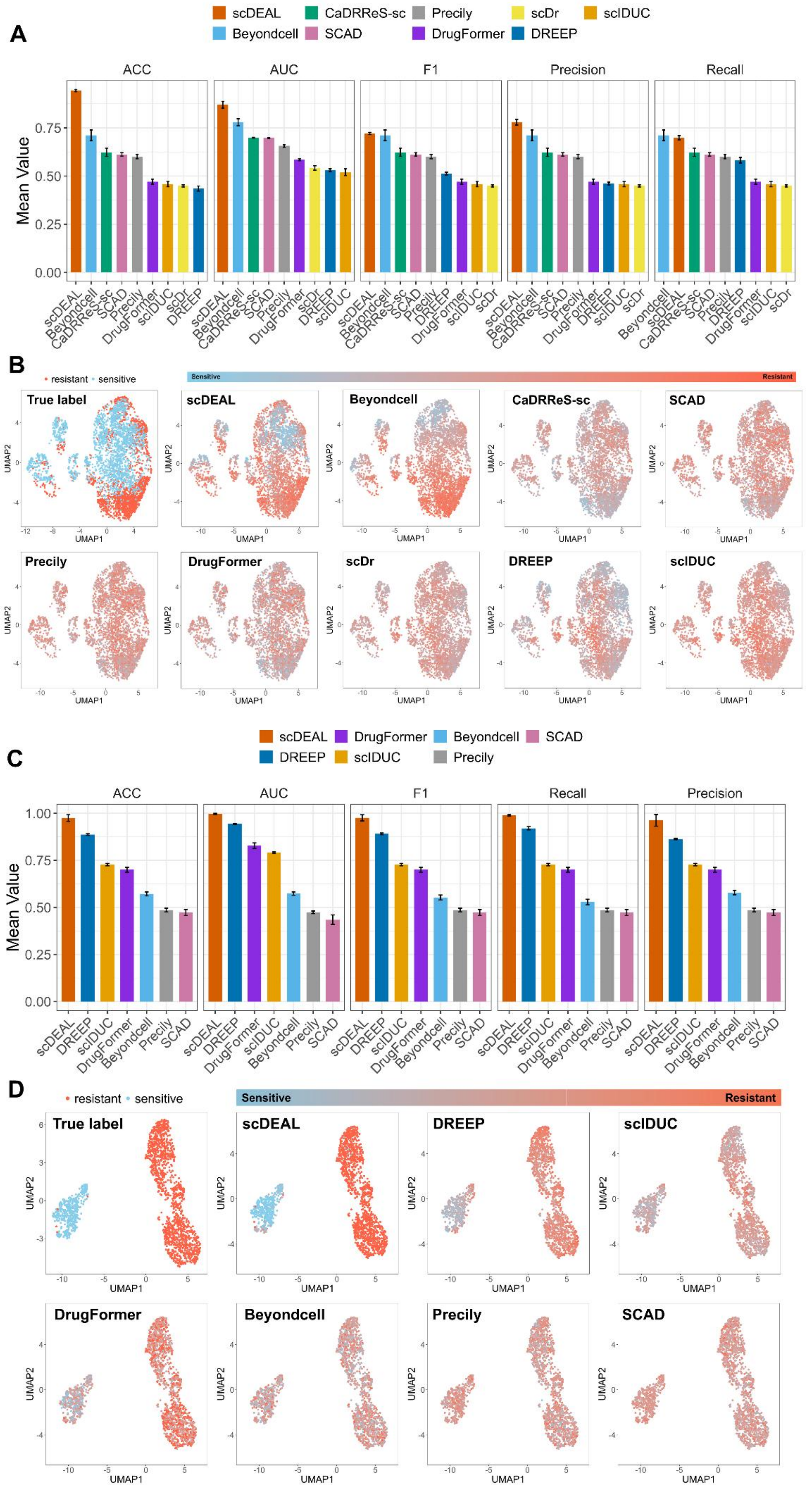
A
B
C
D
scDEAL
Beyondcell
CaDRReS-sc
SCAD
Precily
DrugFormer
scDr
DREEP
scIDUC
ACC
AUC
F1
Precision
Recall
Mean Value
resistant
sensitive
Sensitive
Resistant
True label
UMAP1
UMAP2

**Figure S13. Performance on lineage-tracing single-cell drug response data across treatment states. (A, B)** Results on the ReSisTrace dataset for distinguishing pre-treatment and post-treatment cell populations. **(A)** Mean accuracy, AUROC, F1 score, precision, and recall across all evaluated methods. **(B)** UMAP visualization of ground truth labels and model-predicted scores. **(C, D)** Corresponding results on the Watermelon dataset. **(C)** Mean accuracy, AUROC, F1 score, precision, and recall across all evaluated methods. **(D)** UMAP visualization of ground truth labels and model-predicted scores.

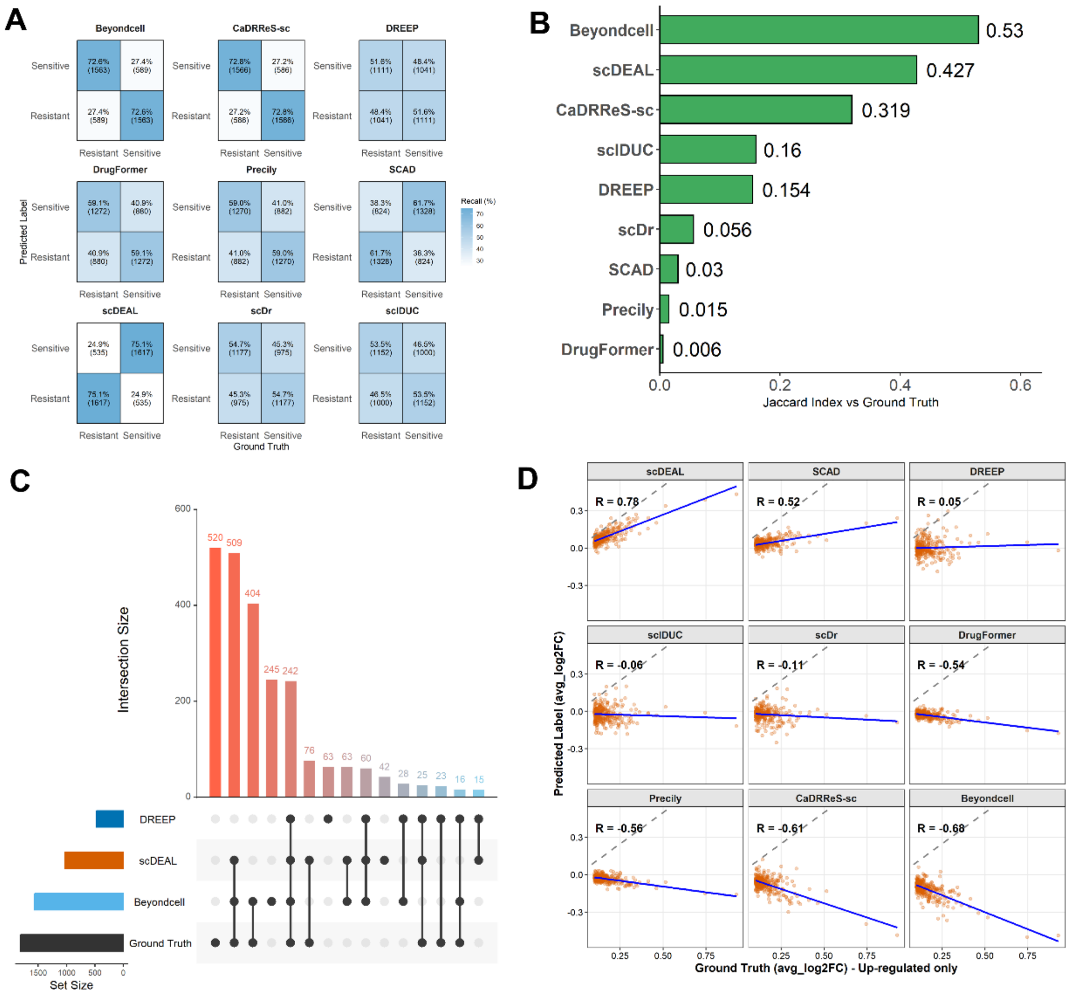


**Figure S14. Prediction concordance analysis between sensitive (pre-treatment) and resistant (post-treatment) cells. (A)** Confusion matrix-based comparison of predicted and ground-truth drug response labels across methods. **(B)** Jaccard-based overlap between DEGs derived from predicted labels and those derived from ground-truth labels. **(C)** Shared and method-specific DEGs derived from predicted labels by the top three methods ranked by Jaccard index and from ground-truth labels. **(D)** Concordance between predicted and ground-truth log2 fold-change profiles.

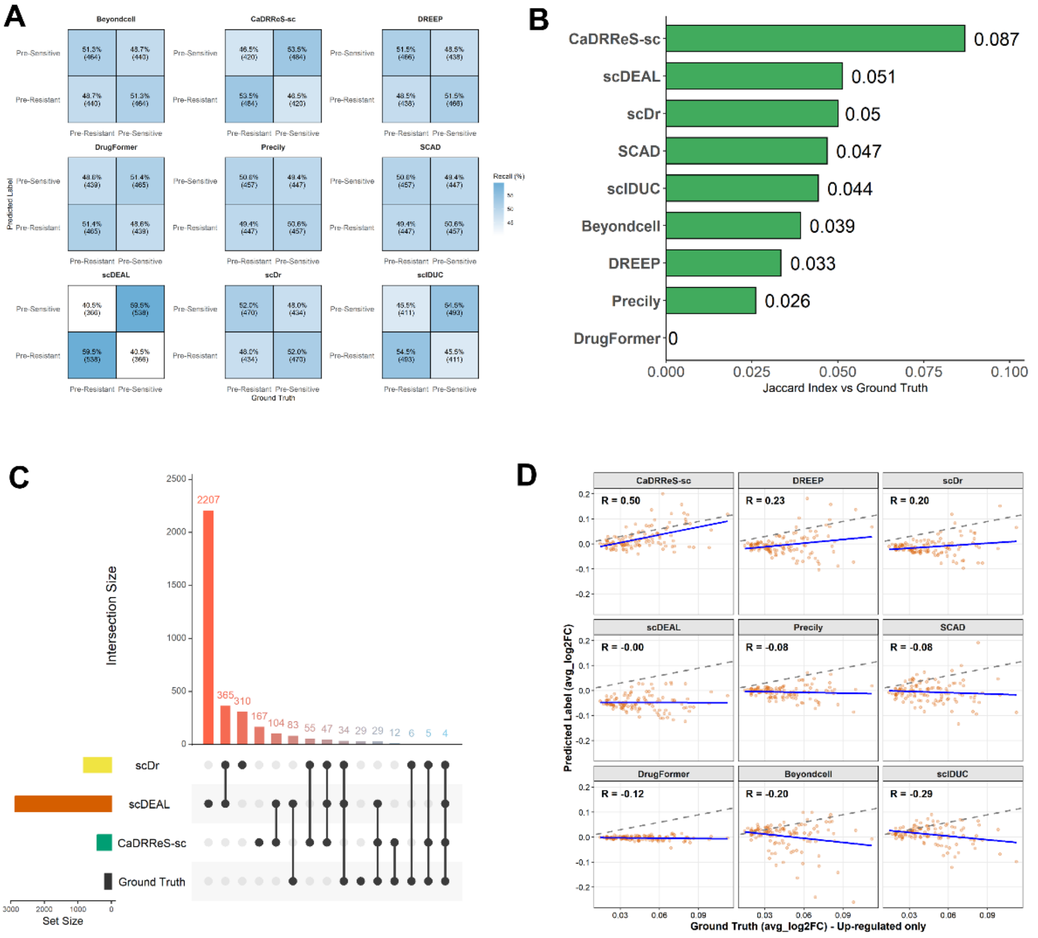


**Figure S15. Prediction concordance analysis between pre-sensitive and pre-resistant cells. (A)** Confusion matrix-based comparison of predicted and ground-truth drug response labels across methods. **(B)** Jaccard-based overlap between DEGs derived from predicted labels and those derived from ground-truth labels. **(C)** Shared and method-specific DEGs derived from predicted labels by the top three methods ranked by Jaccard index and from ground-truth labels. **(D)** Concordance between predicted and ground-truth log2 fold-change profiles.

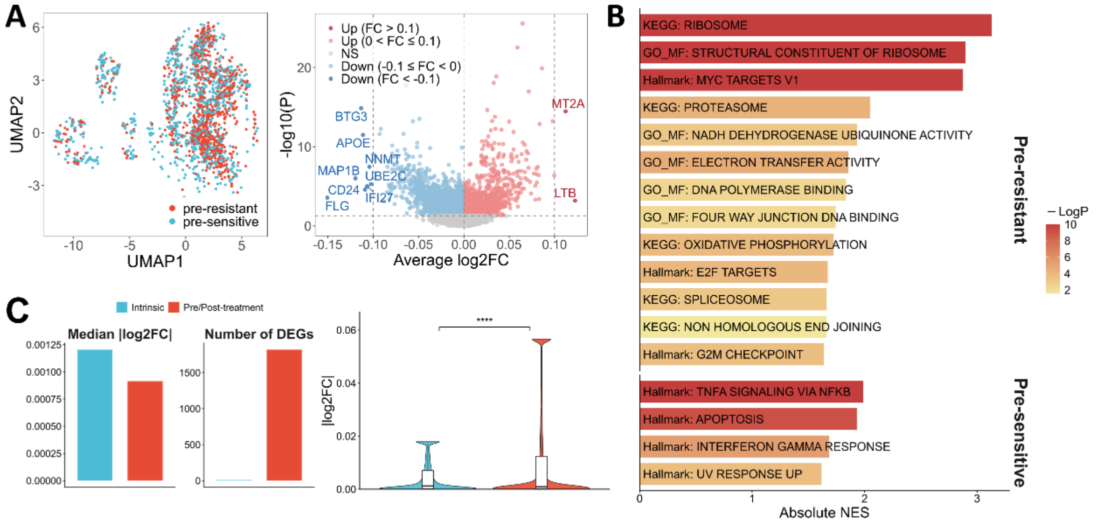


**Figure S16. Transcriptomic characterization of pre-sensitive and pre-resistant cells in the ReSisTrace dataset. (A)** UMAP visualization of pre-sensitive and pre-resistant cells and volcano plot showing differential gene expression between the two populations. **(B)** Pathway enrichment analysis of pre-resistant and pre-sensitive cells using KEGG, GO-MF, and Hallmark gene sets. **(C)** Comparison of differential expression between the pre-sensitive versus pre-resistant and pre-treatment versus post-treatment contrasts, including effect size, DEG number, and absolute log2 fold-change distributions. ****P < 0.0001.

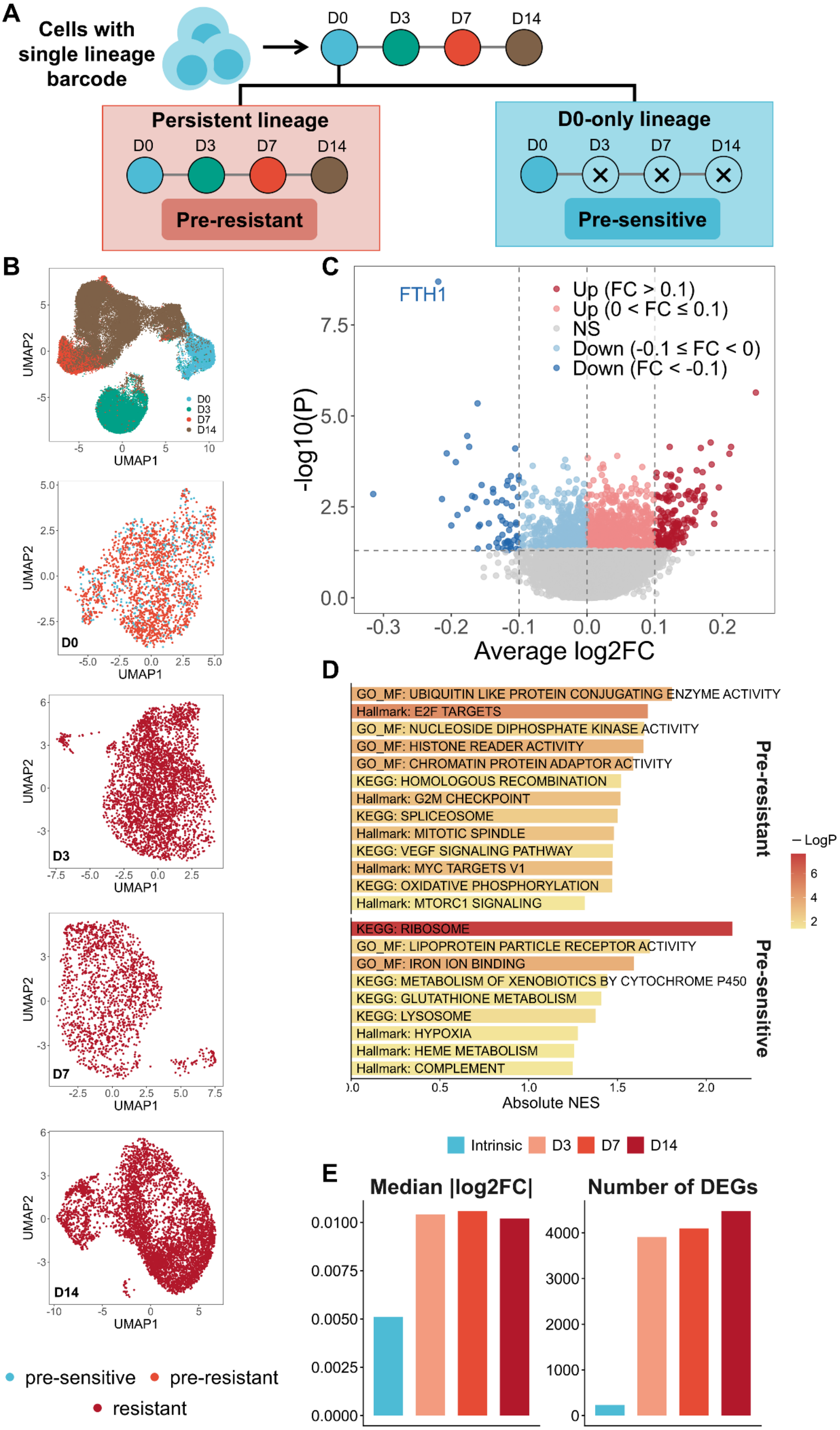
A
Cells with single lineage barcode
D0 D3 D7 D14
Persistent lineage
Pre-resistant
D0-only lineage
Pre-sensitive
B
UMAP1
UMAP2
D0
D3
D7
D14
pre-sensitive
pre-resistant
resistant
C
FTH1
Up (FC > 0.1)
Up (0 < FC ≤ 0.1)
NS
Down (-0.1 ≤ FC < 0)
Down (FC < -0.1)
-log10(P)
Average log2FC
D
GO_MF: UBIQUITIN LIKE PROTEIN CONJUGATING ENZYME ACTIVITY
Hallmark: E2F TARGETS
GO_MF: NUCLEOSIDE DIPHOSPHATE KINASE ACTIVITY
GO_MF: HISTONE READER ACTIVITY
GO_MF: CHROMATIN PROTEIN ADAPTOR ACTIVITY
KEGG: HOMOLOGOUS RECOMBINATION
Hallmark: G2M CHECKPOINT
KEGG: SPLICEOSOME
Hallmark: MITOTIC SPINDLE
KEGG: VEGF SIGNALING PATHWAY
Hallmark: MYC TARGETS V1
KEGG: OXIDATIVE PHOSPHORYLATION
Hallmark: MTORC1 SIGNALING
KEGG: RIBOSOME
GO_MF: LIPOPROTEIN PARTICLE RECEPTOR ACTIVITY
GO_MF: IRON ION BINDING
KEGG: METABOLISM OF XENOBIOTICS BY CYTOCHROME P450
KEGG: GLUTATHIONE METABOLISM
KEGG: LYSOSOME
Hallmark: HYPOXIA
Hallmark: HEME METABOLISM
Hallmark: COMPLEMENT
Pre-resistant
Pre-sensitive
LogP
Absolute NES
E
Intrinsic D3 D7 D14
Median |log2FC|
Number of DEGs

**Figure S17. Transcriptomic characterization of pre-sensitive and pre-resistant cells in the Watermelon dataset. (A)** Schematic illustration of lineage-based definitions of pre-resistant and pre-sensitive cells using longitudinal barcode information from D0 to D14. **(B)** UMAP visualization of cells across time points and of pre-sensitive, pre-resistant, and resistant populations. **(C)** Volcano plot showing differential gene expression between pre-sensitive and pre-resistant cells. **(D)** Pathway enrichment analysis of pre-resistant and pre-sensitive cells using KEGG, GO-MF, and Hallmark gene sets. **(E)** Comparison of differential expression between the intrinsic pre-sensitive versus pre-resistant contrast and the D3, D7, and D14 treatment-associated contrasts, including median absolute log2 fold change and number of differentially expressed genes.

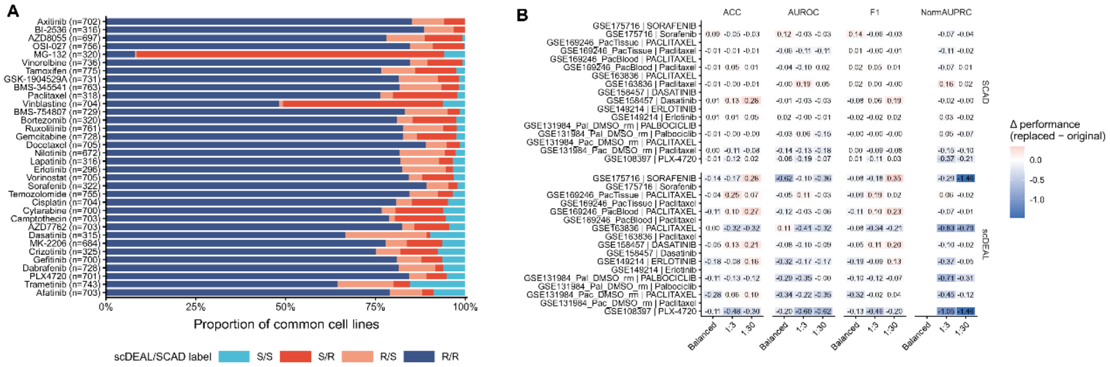


**Figure S18. Dataset-level comparison of training-label agreement and substitution effects. (A)** Proportions of the four scDEAL/SCAD label combinations across shared drugs. **(B)** Dataset-level changes in metrics after training-label substitution under the SCAD and scDEAL architectures across balanced and imbalanced settings.

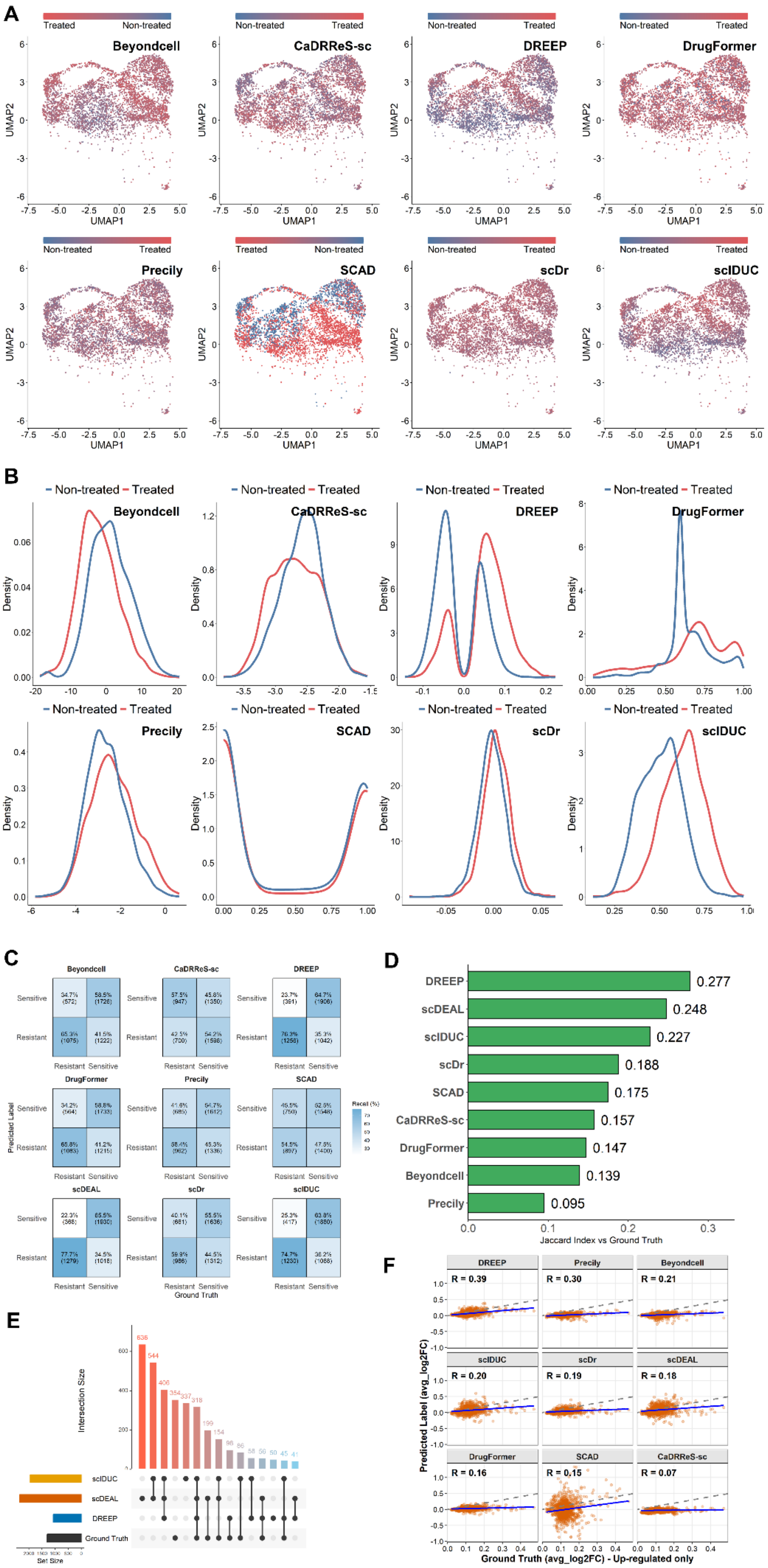
A
Beyondcell
CaDRReS-sc
DREEP
DrugFormer
Precily
SCAD
scDr
scIDUC
Treated
Non-treated
UMAP1
UMAP2
B
Non-treated
Treated
Density
C
Predicted Label
Ground Truth
Sensitive
Resistant
Recall (%)
scDEAL
D
DREEP 0.277
scDEAL 0.248
scIDUC 0.227
scDr 0.188
SCAD 0.175
CaDRReS-sc 0.157
DrugFormer 0.147
Beyondcell 0.139
Precily 0.095
Jaccard Index vs Ground Truth
E
Intersection Size
Set Size
scIDUC
scDEAL
DREEP
Ground Truth
F
R = 0.39
R = 0.30
R = 0.21
R = 0.20
R = 0.19
R = 0.18
R = 0.16
R = 0.15
R = 0.07
Predicted Label (avg_log2FC)
Ground Truth (avg_log2FC) - Up-regulated only

**Figure S19. Predicted score distribution and prediction concordance analysis for the paclitaxel-treated group. (A-B)** Predicted drug response scores shown by UMAP **(A)** and density plots **(B)**. **(C)** Confusion matrix-based comparison of predicted and ground-truth drug response labels across methods. **(D)** Jaccard-based overlap between DEGs derived from predicted labels and those derived from ground-truth labels. **(E)** Shared and method-specific DEGs derived from predicted labels by the top three methods ranked by Jaccard index and from ground-truth labels. **(F)** Concordance between predicted and ground-truth log2 fold-change profiles.

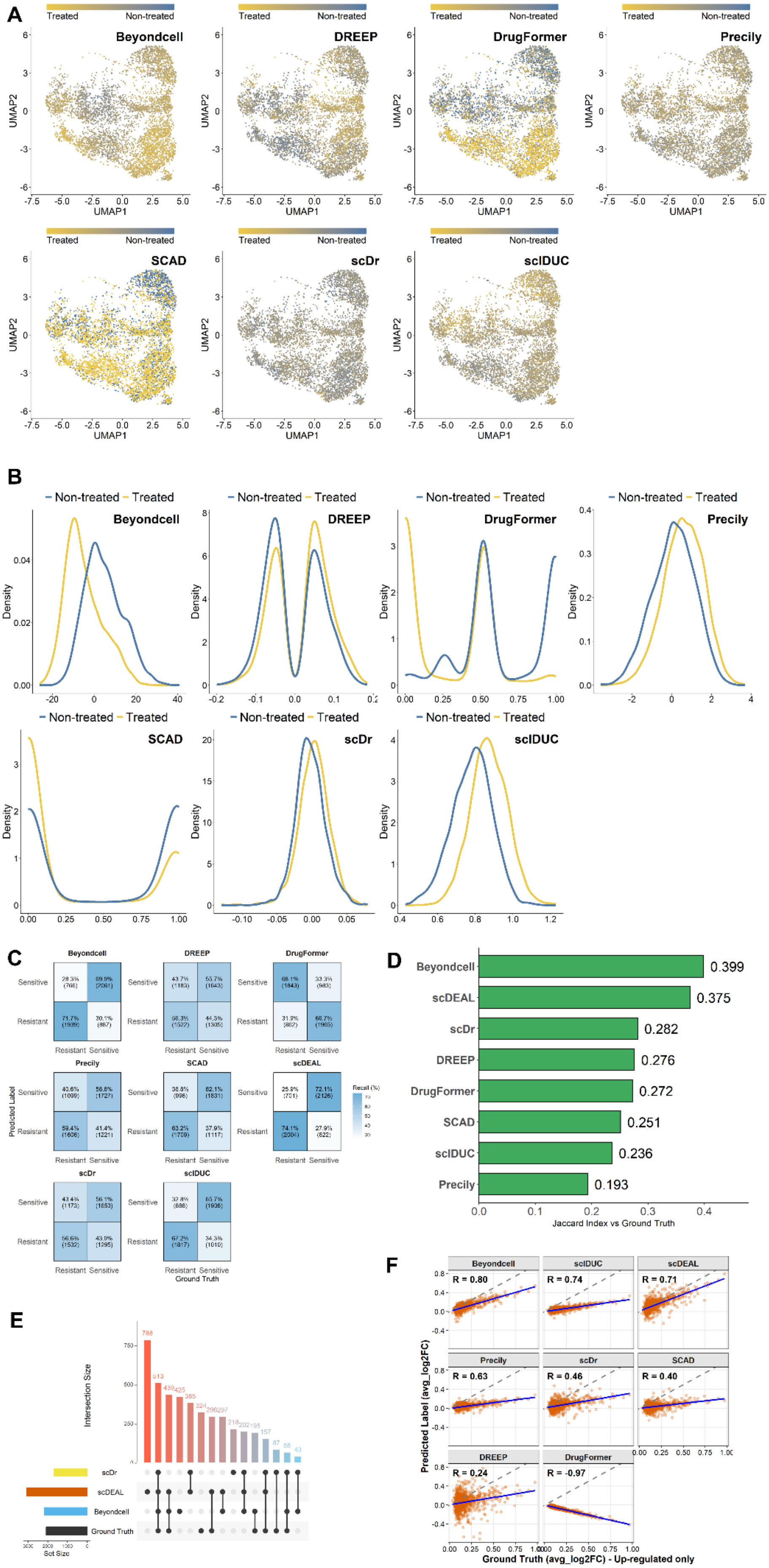
A
Treated
Non-treated
Beyondcell
DREEP
DrugFormer
Precily
SCAD
scDr
scIDUC
UMAP1
UMAP2
B
Non-treated
Treated
Density
C
Sensitive
Resistant
Predicted Label
Ground Truth
Recall (%)
scDEAL
D
Beyondcell 0.399
scDEAL 0.375
scDr 0.282
DREEP 0.276
DrugFormer 0.272
SCAD 0.251
scIDUC 0.236
Precily 0.193
Jaccard Index vs Ground Truth
E
Intersection Size
Set Size
Ground Truth
F
R = 0.80
R = 0.74
R = 0.71
R = 0.63
R = 0.46
R = 0.40
R = 0.24
R = -0.97
Predicted Label (avg_log2FC)
Ground Truth (avg_log2FC) - Up-regulated only

**Figure S20. Predicted score distribution and prediction concordance analysis for the trametinib-treated group. (A-B)** Predicted drug response scores shown by UMAP **(A)** and density plots **(B)**. **(C)** Confusion matrix-based comparison of predicted and ground-truth drug response labels across methods. **(D)** Jaccard-based overlap between DEGs derived from predicted labels and those derived from ground-truth labels. **(E)** Shared and method-specific DEGs derived from predicted labels by the top three methods ranked by Jaccard index and from ground-truth labels. **(F)** Concordance between predicted and ground-truth log2 fold-change profiles.

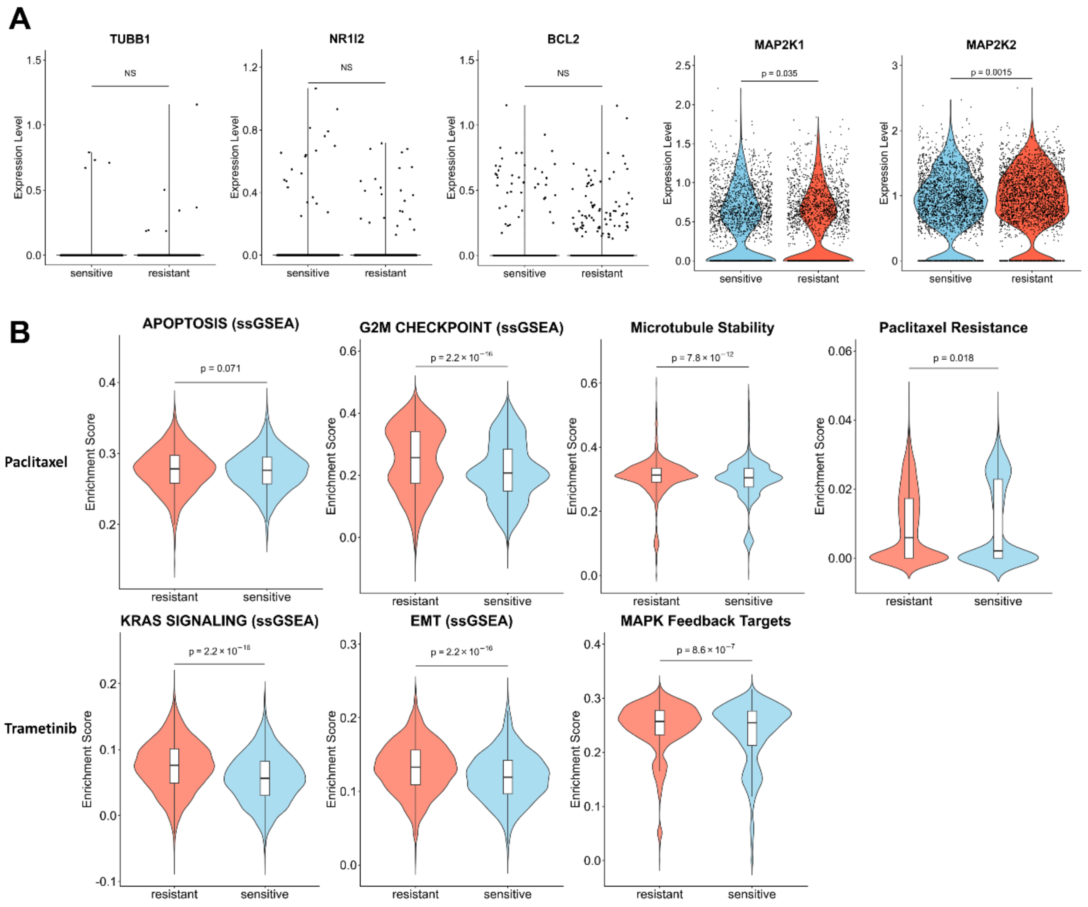


**Figure S21. Mechanistic validation of scDEAL predictions. (A)** Target gene expression in predicted sensitive versus resistant cells. NS: non-significant; Wilcoxon rank-sum test. **(C)** ssGSEA scores for drug-specific pathways and resistance signatures across predicted groups. P-values were calculated using the Wilcoxon rank-sum test.

**References**


1. Collins, R.L., et al., *A cross-disorder dosage sensitivity map of the human genome.* Cell, 2022. **185**(16): p. 3041-3055 e25.
2. Makinen, L., et al., *Pancreatic Cancer Organoids in the Field of Precision Medicine: A Review of Literature and Experience on Drug Sensitivity Testing with Multiple Readouts and Synergy Scoring.* Cancers (Basel), 2022. **14**(3).
3. Boj, S.F., et al., *Organoid models of human and mouse ductal pancreatic cancer.* Cell, 2015. **160**(1-2): p. 324-38.
4. Hao, Y., et al., *Integrated analysis of multimodal single-cell data.* Cell, 2021. **184**(13): p. 3573-3587 e29.
5. Korsunsky, I., et al., *Fast, sensitive and accurate integration of single-cell data with Harmony.* Nat Methods, 2019. **16**(12): p. 1289-1296.
6. Hanzelmann, S., R. Castelo, and J. Guinney, *GSVA: gene set variation analysis for microarray and RNA-seq data.* BMC Bioinformatics, 2013. **14**: p. 7.
7. Aibar, S., et al., *SCENIC: single-cell regulatory network inference and clustering.* Nat Methods, 2017. **14**(11): p. 1083-1086.
8. Qiu, X., et al., *Single-cell mRNA quantification and differential analysis with Census.* Nat Methods, 2017. **14**(3): p. 309-315.
9. Zhang, J., et al., *ClusterGVis: An Advanced Visualization and Clustering Tool for Gene Expression Analysis.* Genomics Proteomics Bioinformatics, 2026.
10. Subramanian, A., et al., *Gene set enrichment analysis: a knowledge-based approach for interpreting genome-wide expression profiles.* Proc Natl Acad Sci U S A, 2005. **102**(43): p. 15545-50.
11. Liberzon, A., et al., *The Molecular Signatures Database (MSigDB) hallmark gene set collection.* Cell Syst, 2015. **1**(6): p. 417-425.
12. Wu, T., et al., *clusterProfiler 4.0: A universal enrichment tool for interpreting omics data.* Innovation (Camb), 2021. **2**(3): p. 100141.
